# Serotype swapping in *Klebsiella* spp. by plug-and-play

**DOI:** 10.1101/2025.08.29.672822

**Authors:** Julie Le Bris, Hugo Varet, Eduardo P.C. Rocha, Olaya Rendueles

## Abstract

Understanding how complex, multi-gene systems evolve and function across genetic backgrounds is a central question in molecular evolution. While such systems often impose costs through epistatic interactions, some may behave as modular, “plug-and-play” units that retain function with minimal disruption.

We used the polysaccharide capsule locus of *Klebsiella pneumoniae*, a highly exchangeable and fast-evolving locus, as a model. We genetically engineered capsule exchanges (swaps) across diverse genetic backgrounds and combined transcriptomics, fitness assays, and evolution experiments, to show that capsule exchange has negligible effects on global expression and only marginal fitness costs, regardless of serotype. Adaptation to capsule-costly environments consistently reduced capsule production, regardless of serotype, revealing shared adaptive trajectories rather than serotype-specific pathways. Moreover, serotype-specific traits involved in bacterial virulence, such as biofilm formation and hypermucoviscosity, were conserved across genetic backgrounds. This reveals that capsule swapping can directly shape host-pathogen interactions and influence within-patient evolution.

Our findings provide strong evidence that capsule loci display plug-and-play dynamics: they are transferable, functional across contexts, and minimally disruptive to the host genome. This allows capsules to be seamlessly swapped, and help explain the evolutionary success, ecological versatility, and pervasive exchangeability of capsules in *K. pneumoniae*.

## INTRODUCTION

Bacteria continuously expand their functional repertoire through a variety of evolutionary mechanisms, including gene duplication and diversification^1,2^, domain rearrangement, and horizontal gene transfer. These processes can result in acquisition of analogous functions or in novel ones, amongst which, antibiotic resistance, virulence, or metabolic pathways ^3,4^. While numerous studies have documented the advantages of such gains in the laboratory and natural populations, evolutionary success is mostly rare^5,6^. The integration of such systems into existing cellular networks may not be seamless and can come at a fitness cost due to regulatory conflicts, metabolic imbalances, or structural incompatibilities^7^. Their success depends on the gene-by-environment fitness effect^8^ and on complex epistatic interactions with the host genome^9^, where the impact of a gene is shaped by the recipient genetic background. Indeed, epistasis is widespread in nature, influencing evolutionary dynamics in cancers, viruses^10^, and bacteria^11,12^. It can be positive, for instance, large plasmids can promote the maintenance of other, smaller plasmids in bacterial populations^13^, but it can also be negative^12,14^, requiring compensatory mutations to limit the fitness costs^15,16^. These interactions underscore the context-dependency of genetic innovation and the challenges of functional integration.

Promiscuous systems which are frequently mobilized across genetic backgrounds are hypothesized to follow a plug-and-play evolutionary dynamic, whereby successful integration results in minimal epistatic interference with the host genome^17,18^. Such systems are characterized by a high degree of modularity, that is, their functionality is largely self-contained and does not rely on tight interactions with host-specific pathways. This architecture allows expression and maintenance of the novel function with limited physiological burden or need for compensatory mutations. The plug-and-play model has been extensively documented in phage biology and protein domain shuffling^19,20^, where modular elements maintain functionality across hosts. Modularity enhances evolvability by enabling the reconfiguration of cellular functions through the acquisition or exchange of discrete, low-conflict genetic modules in response to changing environmental conditions^21,22^. Despite its conceptual appeal, whether true plug-and-play dynamics apply to large, multi-gene chromosomal systems in bacteria remains untested. Notably, a key question is whether such systems impose universal fitness trade-offs upon transfer or whether their architecture allows for broad functional compatibility with minimal host rewiring.

An example of a complex multi-gene system is the extracellular polysaccharide capsule locus (*cps*). It is one of the fastest-evolving loci in Bacteria due to horizontal gene transfer and characterized by elevated recombination rates^23,24^. Present in many facultative pathogens and widespread in environmental species^25^, this highly diverse surface structure is a major virulence factor with critical roles in microbial ecology and evolution. Its impact further extends to clinical settings, biotechnology and public health^26^. Group I capsules, the most prevalent, also known as Wzy/Wzx-dependent capsules, contain both conserved core genes and a variable region encoding oligosaccharide modifying and polymerization enzymes. This variable region results in different biochemical compositions, also known as serotypes^27,28^. Capsule locus diversity is shaped by relaxed purifying selection combined with strong diversifying selection^27,28^, likely driven by the immune system, phage and/or protist predation^26–28^.

In the enterobacteria *Klebsiella pneumoniae* (*Kpn*), an opportunistic pathogen able to colonize a broad range of environments^29,30^, specific capsule types have been linked to distinct diseases, such as K3 with rhinoscleromatosis^31^ and K2 with inflammatory bowel diseases^32,33^. Recent findings suggest that serotype influences *Kpn* bloodstream survival rates^34^, pointing to serotype-specific contributions to virulence. In *Kpn*, over 140 different serotypes have been described^35^. The current model posits that the whole capsule locus is transferred horizontally, most likely by conjugative elements. Once in the recipient cell, the capsule integrates in a one-step large recombination event spanning the whole conserved locus known as the K-locus (10-30 kb)^24^. These exchanges occur preferentially across genomes encoding biochemically similar capsules, rather than by genetic relatedness thereby suggesting important epistatic effects between host genome and the K-locus^24^. This presents an interesting evolutionary conundrum, as the pervasiveness of capsule exchanges suggests this process occurs by a plug-and-play process in which there is minimal disruption to the cell, e.g. low negative epistasis. Further, the conserved core genes of the locus are more integrated in the cell metabolism, and their exchange may have a stronger impact on host fitness^36^, whereas the non-core (accessory), serotype-specific genes, are expected to have low levels of negative epistasis as they have not co-evolved with the host genome. Finally, capsule exchanges also reshape the surface glycobiology, potentially altering electrostatic properties and disrupting cellular homeostasis.

Several studies have engineered *in vitro* capsule locus swaps in *Kpn*, primarily to address the inheritance of virulence alongside capsule serotypes^34,37,38^. Yet, the broader evolutionary consequences on capsule exchange, such as its effect on bacterial fitness and physiology remain unexplored. We hypothesized that if there were any transcriptional changes or fitness costs, these should occur in capsule exchanges between the most biochemically different serotypes. To test this, we expanded a collection of strains in which different *cps* loci were introduced in different genetic backgrounds^39^, and leveraged transcriptomic profiling with large-scale pairwise competitions and evolution experiments. This integrative framework allowed us to directly quantify both regulatory and fitness consequences of serotype exchange, and compare the effects across serotypes at different steps of the infectious process. Our results show that there is little if no impact in the regulatory network of the cell and marginal metabolic or energetic burden. Our study provides a clear example of plug-and-play in Bacteria and demonstrates that the widespread transferability of the *cps* locus across *Klebsiella* species and beyond is underpinned by true modularity.

## RESULTS

### Genetic background is the primary determinant of capsule production

To explore potential fitness effects and epistatic interactions between the capsule locus and its host genome upon integration by horizontal gene transfer, we expanded a collection of strains to encompass five native strains, their five respective acapsular mutants (Δ*cps*) alongside 19 capsule-swapped strains. Specifically, five phylogenetically diverse genetic backgrounds – three hypervirulent (*Kpn* NTUH K2044, *Kpn* BJ1 and *Kpn* CIP 52.145), one commensal (*Kpn* ST45) and one environmental (*K. variicola* -*Kva*-OM26) strain (Table 1)– expressed one of four clinically relevant and well-characterized serotypes (Figure S1) associated to hypervirulence (K1 & K2), rhinoscleromatitis (K3)^31^, and carbapenem-resistance (K24). Of note, K3 could not be introduced into a *Kva* OM26 genetic background. These 19 capsule-swapped strains include the control strains whereby each strain’s native capsule serotype was reintroduced into the Δ*cps* mutant. In two genetic backgrounds (*Kpn* NTUH K2044 and *Kpn* CIP 52.145), complementation involved an alternative capsule locus of the same serotype, which exhibited minor genomic differences, and resulted in minor, but significant, differences in capsule production (Table 1, Figure S1, Figure S2A). These strains will be collectively referred to as capsule-swapped strains. Their genetic and biochemical similarities are portrayed in Figure 1.

**Figure 1.**
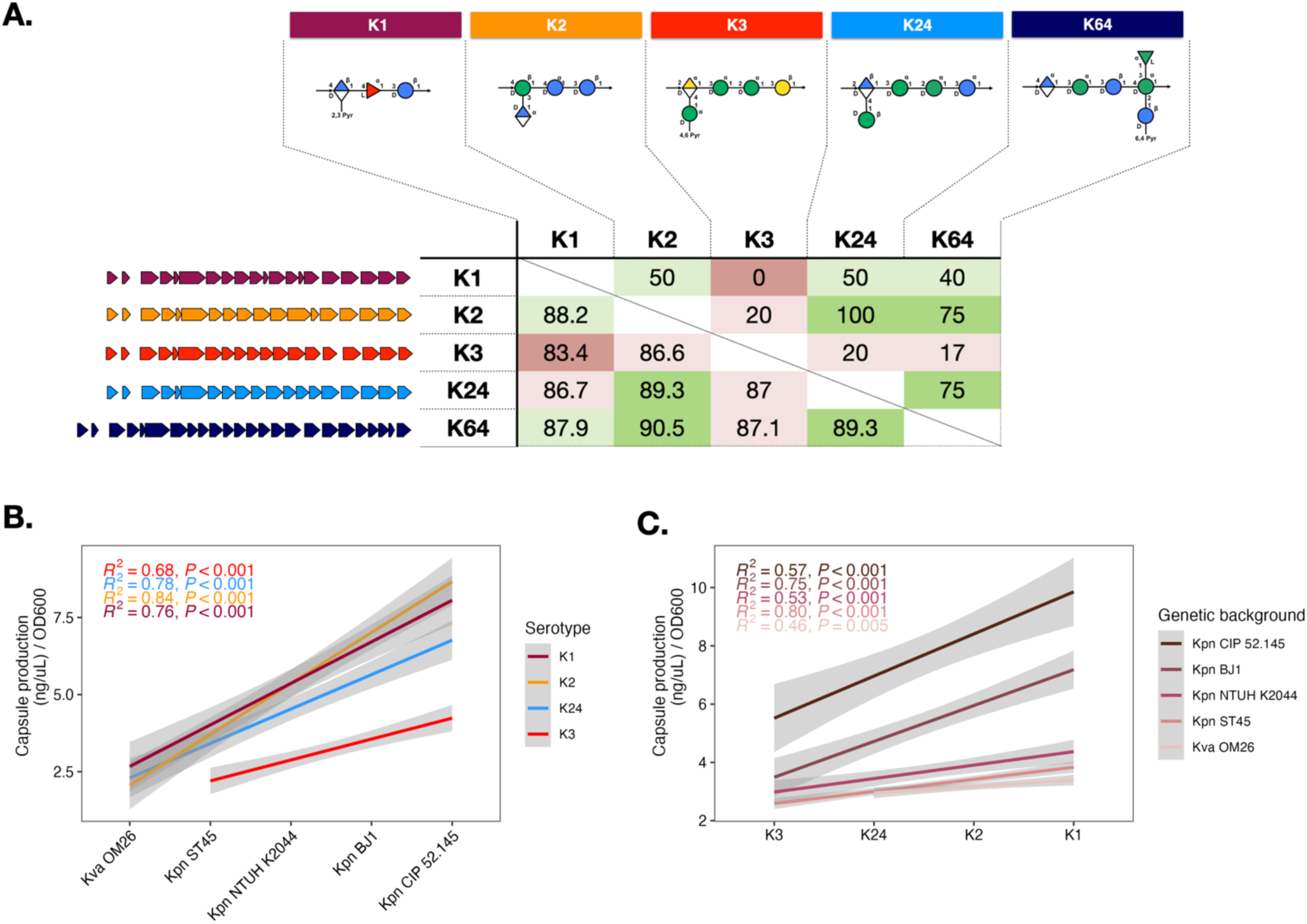
Characteristics and capsule production of capsule-swapped strains. **A.** Genetic (lower triangle) and biochemical relatedness (upper triangle) between the different capsule serotypes analyzed in this study. Biochemical similarity was calculated as the percentage of residues shared between two serotypes (# of shared unique oligosaccharides / # of different non-redundant oligosaccharides). Oligosaccharidic compositions and models were taken from KPAM^42^. Genetic relatedness was calculated taking into account all shared genes between two serotypes (protein by protein alignment, bidirectional identity averaged by pairwise comparison across proteins). **B-C.** Capsule production of each capsule-swapped strain determined by the uronic acid method. The genetic background (B) or serotype (C) is indicated on the x-axis. Different colored lines represent linear regressions for each serotype (B) or genetic background (C) (R^2^). Capsule production of the native strains is shown in Figure S2A and each strain detailed in Figure S2B-C.

**Table 1.** Genetic backgrounds and serotypes used in this study to generate all the capsule-swapped strains. The identities of the different native capsule loci compared to those reintroduced as controls are included.

| **NATIVE STRAINS** |  |  |  |  |
| --- | --- | --- | --- | --- |
| **Strain** | **Native serotype** | **O-antigen type** | **Sequence Type** | **Origin** |
| *Kpn* NTUH K2044 | K1 | O1$\alpha\beta$,2β | ST23 | Human liver abscess, Taiwan<sup>40</sup> |
| *Kpn* BJ1 | K2 | O1$\alpha\beta$,2α | ST380 | Human liver abscess, France<sup>41</sup> |
| *Kpn* CIP 52.145 | K2 | O1$\alpha\beta$,2α | ST66 | Human liver abscess, Indonesia<sup>41</sup> |
| *Kpn* ST45 | K24 | O2α | ST45 | Human carriage, feces, Netherlands<sup>41</sup> |
| *K. variicola* OM26 | K64 | O3$\alpha\beta$ | | Environmental, France, CIP 80.47 |
| **SEROTYPES** |  |  |  |  |
| **Capsule locus** | **Strain of origin** | **Other variants & % of identity** |  |  |
| K1 | *Kpn* SA12 | K1_NTUH K2044; 99.98% identity (SNPs including in *wzc*) |  |  |
| K2 | *Kpn* BJ1 | K2_CIP 52.145; 98.24% identity (truncated *oatWY*) |  |  |
| K3 | *Kpn* ATCC13883T |  |  |  |
| K24 | *Kpn* ST45 |  |  |  |

We first verified that all capsule-swapped strains produced detectable levels of extracellular capsule (Figure 1B-C), and that serotype complementation would result in similar levels of capsule production (Figure S2A). Capsule quantification revealed that the native serotype was not necessarily the most expressed in the original genetic background (Figure 1C), and that different strains produce different amounts of capsule (Figure 1B, multifactorial ANOVA, p< 2e-16, Table S1).

In addition to the three well-known promoters that directly regulate the capsule locus^43^ (Figure S1), ca. one hundred recently identified capsule regulators are located elsewhere in the genome^44,45^, raising the question of whether capsule production is driven more by the genetic background or the serotype. A stepwise linear regression model reveal that the genetic background is the main factor explaining variation in capsule production. Interestingly, across all serotypes, capsule production followed the same trend: *Kpn* CIP 52.145 > *Kpn* BJ1 > *Kpn* NTUH K2044 > *Kpn* ST45 > *Kva* OM26 (Figure 1B, Figure S2B). Also, independently of the genetic background, serotypes were associated with different levels of capsule production in the ranking order: K1 > K2 > K24 > K3 (Figure 1C, Figure S2C). Our data show that capsule production follows a strong hierarchical pattern dictated by the genetic background and the serotype. Specifically, certain genetic backgrounds consistently produce a higher level of capsule regardless of serotype, while some serotypes are similarly expressed across diverse genetic contexts. Further, the interaction between genetic background and serotype in our regression model resulted in a low F-value (F = 7.9), indicating limited interplay between these two factors. Collectively, our data suggests that intrinsic properties of capsule loci are conserved among strains and that the variance observed is mostly explained by the serotype and the genetic background.

### Newly acquired capsule loci result in minimal gene expression changes

The adaptive response to a novel genetic element or function can occur either through transcriptional changes or genetic modifications. While the latter are difficult to revert^46,47^, transcriptional response is quick, involves no lasting costs, and is not passed on to future generations^48^. We hypothesized that new serotypes could alter surface properties –causing protein jamming, electrostatic shifts, and membrane disorganization– especially with decreasing biochemical relatedness between the native serotype and the new one (Figure 1A). Also, we expected that gene expression changes following capsule exchange will be more similar between serotypes with greater biochemical relatedness. To study this, we performed a RNAseq analysis of capsule-swapped strains during exponential phase in nutrient-poor medium where the capsule is produced at higher levels^30^. Acapsulated mutants of each genetic background were also sequenced and used as controls (see Methods).

We first investigated the expression of the different capsule serotypes across the different backgrounds compared to their native capsule (see Methods). Among the 194 capsule genes analyzed, across all serotype and genetic background combinations, only 18 genes were differentially expressed (nine upregulated and nine downregulated) (Figure S3). We performed a Principal Component Analyses (PCA) (see Methods), to test if each serotype is associated with a specific transcriptional profile when expressed in different genetic backgrounds. Capsule serotypes are separated by the first two components, although only very mildly so, with the exception of K3 (Figure S4A-B). This indicates that the gene expression pattern of K3 capsule is different from the others. Indeed, K3 is the most distinct serotype in the set, sharing the fewest number of sugar residues with other serotypes (either one or none) (Figure 1A). At the gene level, very few commonalities in terms of gene expression patterns are observed, if any. In *Kpn* ST45, the first enzyme of the capsule biosynthesis pathway (*wcaJ*) is significantly downregulated upon introduction of any new serotype (Figure 2A). Yet, the same was not observed in other genetic backgrounds. Further, some genes like *wza*, *wzb* and *wzc* are downregulated in ST45-K3 whereas *wza* is upregulated in OM26-K1.

**Figure 2.**
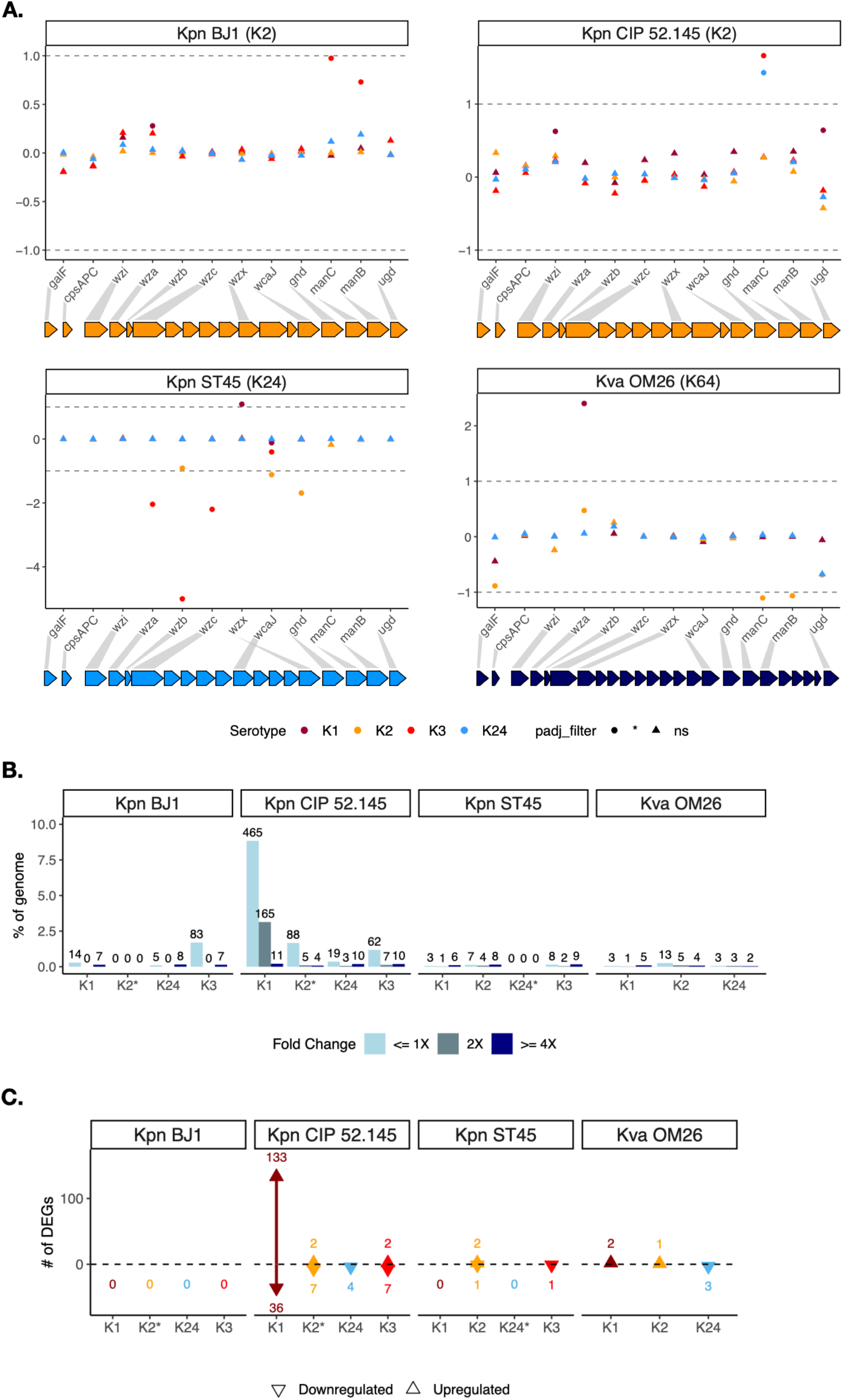
Newly acquired capsule loci result in minimal gene expression changes. **A.** Log_2_FC (fold change) of core capsule locus genes, i.e. genes common to all the serotypes. The shape indicates the significance of the adjusted p-value and the color represents the capsule serotype. The genetic background is indicated at the top of each subpanel. **B.** Percentage of differentially expressed genes (DEGs, adjusted *P* < 0.05) in each genetic background when considering all genes including the capsule locus. DEGs are subdivided according to their respective Fold Change indicated by the color. Corresponding raw number of DEGs is indicated at the top of the bar. **C.** Number of DEGs (adjusted *P* < 0.05 with |log_2_ fold-change| > 1) per capsule swap in each genetic background, either up or down regulated when considering all genes except the capsule locus. Asterisk beside K loci (*) indicates the native serotype of each strain.

We next tested whether the introduction of a novel serotype affects gene expression beyond the capsule locus. PCA analyses of the first two components did not allow the separation by serotype, indicating that the transcription profiles in a given genetic background were very similar and independent of the expressed serotype (Figure S4C). In most strains, we observe that only a small subset of the gene repertoire (x̅ = 1.4%, median= 0.36%, but reaching 8.8%, depending on the background), has a significantly altered expression, but these changes are marginal in magnitude (adjusted *P* < 0.05 with |log_2_ fold-change| < 1) (Figure 2B). Only very few genes were largely differentially expressed (adjusted *P* < 0.05 with |log_2_ fold-change| > 1) (Figure 2C). Contrary to our expectations, there were no changes in proteins associated to membrane biogenesis or homeostasis or in the core metabolism, or in any other pathway (Figure 2C). Only in ST66, the integration of the K1 capsule locus resulted in a significantly higher number of differentially expressed genes (176 genes) —mostly upregulated—compared to any other combination.

Collectively, our results indicate that capsule expression is fairly constant independently of the genomic context, as suggested by the capsule production quantifications mentioned above. Further, despite the changes on the surface imposed by radically different glycobiology of the different serotypes studied here, our analyses show that introduction of novel serotypes does not result in a major cellular transcriptional rewiring.

### Capsule exchanges impose minimal fitness costs and reveal serotype-dependent transitive fitness hierarchies

Given the few transcriptional changes observed, and their magnitude, we hypothesized that serotype exchanges could follow a plug-and-play dynamic, resulting in low/no detectable fitness burden. To test this, we first measured growth in nutrient-rich (LB) medium, where capsules were shown to be costly, and in nutrient-poor (M02) medium where capsules provide a large fitness advantage^30^. We compared the capsule-swapped strains to the native serotype by calculating the area under the curve (AUC) which takes into account lag time, generation time and maximum yield. Our results show that strains with new serotypes do not grow significantly less, independently of the environment (Figure 3A and Figure S5A). Across all strain x environment growth tests (N=38), in only five instances, capsule-swapped strains grew significantly less, whereas in ten instances they grew significantly better (Figure S5A). Our data reveal a negative correlation between the amount of capsule produced in nutrient rich medium and growth (Figure 3B). Specifically, the replacement of the large K1 capsule by any other serotype resulted in significant growth benefits in rich-medium (Figure S5A). Using a stepwise linear regression model, we show that when the capsule is most costly (LB), both the genetic background first, and then the serotype drive growth (Table S2). Yet when the capsule, irrespective of the serotype, is advantageous, growth was primarily governed by the genetic background. Finally, as observed for capsule production, there is little interaction between serotype and genetic background (Table S1 and S2).

**Figure 3.**
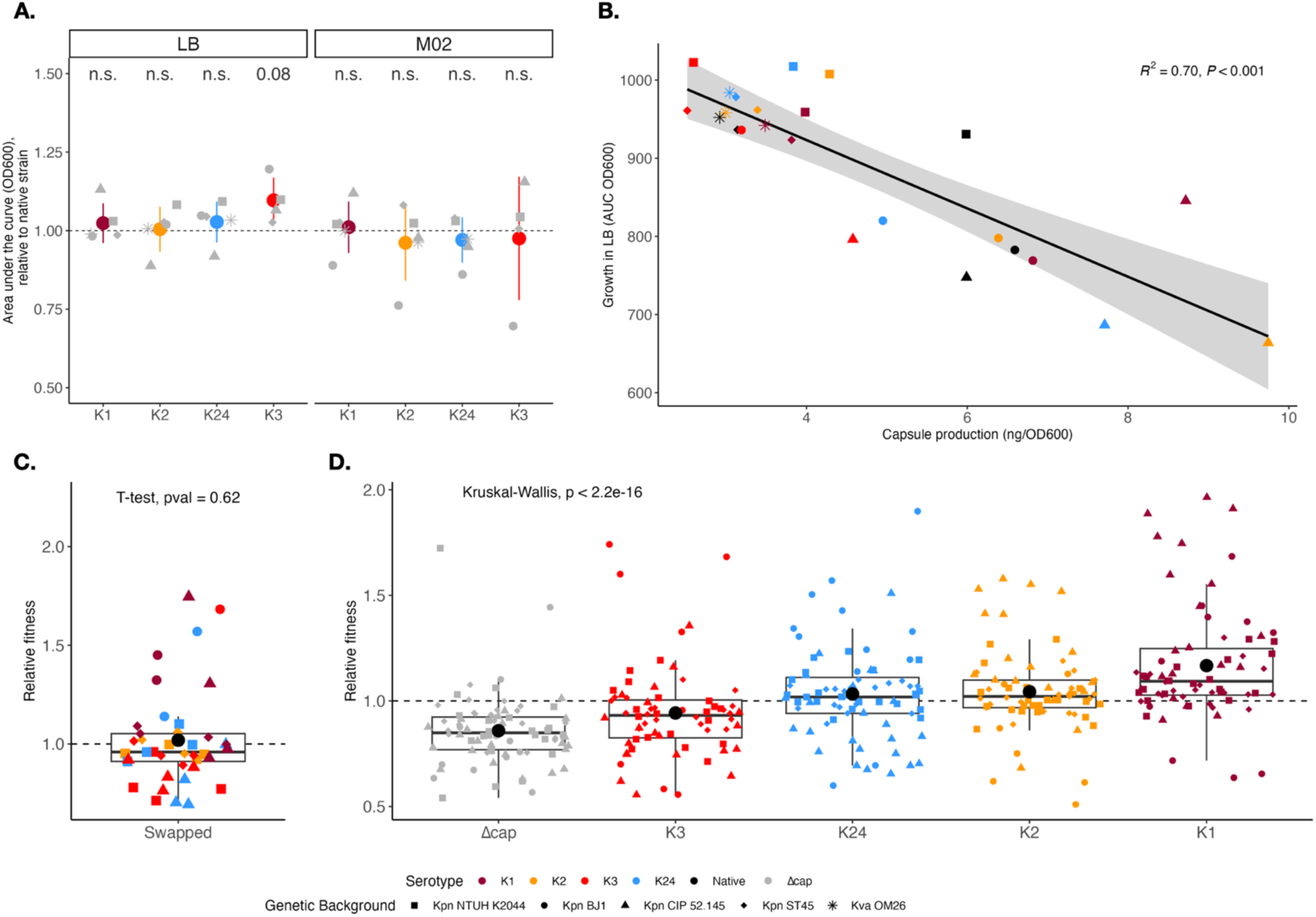
Growth and fitness effect of inserting a novel capsule serotype across genetic backgrounds. **A.** Comparison of the area under the growth curve (AUC OD_600_) between native strains (dashed line) and capsule-swapped strains. AUC was estimated by the formula trapz from the pracma package in R. Each point represents the mean of at least three independent biological replicates. Individual error bars are not shown for visibility purposes. **B.** Correlation between the amount of capsule produced and the growth (area under the curve, AUC) of each capsule-swapped strain. The shape of the points corresponds to the genetic background and color to the serotype. Each point represents the mean of at least three independent biological replicates. Error bars are not included for visibility purposes. Black line represents a linear regression (R2). **C.** Fitness of capsule-swapped strains in direct competition with the native strain, as measured by flow cytometry after 24 hours of coculture. P-value corresponds to Wilcoxon test. **D.** Relative fitness of capsule swap strains grouped by serotype. P-value correspond to Kruskal-Wallis means comparison test.

To further assess the serotype effect on the fitness of the host, we performed fluorescent-based *in vitro* competition assay in nutrient-poor medium. We chromosomally tagged the strains at the attachment site of the transposon Tn7 (attTn7) with red fluorescent (mCherry) protein coding gene. We performed all pairwise competitions in all four *Kpn* genetic backgrounds (NTUH K2044, BJ1, CIP 52.145 and ST45) (N = 336 competitions). We first confirmed that the cost of the fluorescent protein expression was negligeable and that the reinsertion of the native serotype in acapsular mutants did not alter fitness compared to the native strain (Figure S5B). We then tested whether in direct competition with the native serotype, a new serotype had reduced fitness. Surprisingly, we found no significative differences (Figure 3C). More interestingly, our data show that fitness of capsule-swapped strains is mainly dictated by the serotype (multifactorial ANOVA, p<2e-16, Table S1). The relative fitness shows monotonic increased from the acapsular mutant to K1 (Figure 3D). Statistical analysis of the fitness hierarchy using Kendall’s correlation rank confirmed such transitive relation (τ= 0.791, p=5.3644e-6). Further, stepwise linear regression ranked the serotypes as follows: K3 < K24 < K2 < K1 with K1 being the fitter serotype in nutrient-poor medium (Table S2). Such ranking inversely mirrors the cost during growth in nutrient-rich environments, where the capsule is selected against. Of note, a slight cost was observed for *mCherry* integration in the K1 swap of *Kpn* CIP 52.145, the fittest capsule-swapped strain, suggesting its fitness advantage may be underestimated (Figure S5B).

Altogether, our findings reveal that *Klebsiella* capsule-swapped strains fitness is dictated by serotype, and more particularly, by the thickness of the capsule, where in nutrient-poor medium, larger capsules like K1 are fitter than K2, K24 and the thinnest K3^39^.

### Thickness and common genetic pathways drive capsule inactivation of native and novel serotypes

To further corroborate the marginal, if any, cost of introducing a new capsule serotype, we performed a short evolution experiment. Previous research shows that in rich media, acapsular clones rapidly emerge within 20 generations, but in poor media, all clones retain their capsules^30^. If there is a fitness cost associated to the newly acquired capsule, we would expect acapsular mutants to be more strongly selected and emerge faster compared to the native strain. To test this, we performed daily transfers of strains for fifteen days in both nutrient-rich (LB) and -poor (M02) media. This accounts for ca. 100 generations. We visually characterized and counted colonies from all populations and scored the percentage of acapsular clones every day. In line with our previous experiments, in nutrient-poor media, no capsule inactivation was observed regardless of the serotype and the genetic background (Figure S6A).

On the contrary, in nutrient-rich medium, the proportion of capsulated clones rapidly decreased in all the capsule-swapped strains during the first ten days (Figure 4A). However, emergence of acapsular clones was not faster in the swapped strains compared to their native strain, as revealed by the area under the curve of capsule inactivation. A closer inspection of the curves revealed that, during the five first days, capsule inactivation was faster in populations with K1 and K2 serotypes expressing thicker capsules compared to K3 and K24 swaps (Figure 4C, Figure S6B). This is in line with the serotype ranking observed when considering their capsule production and thickness, and inversely mirrors the fitness advantages provided in nutrient-poor medium (Figure 3D)^39^.

**Figure 4.**
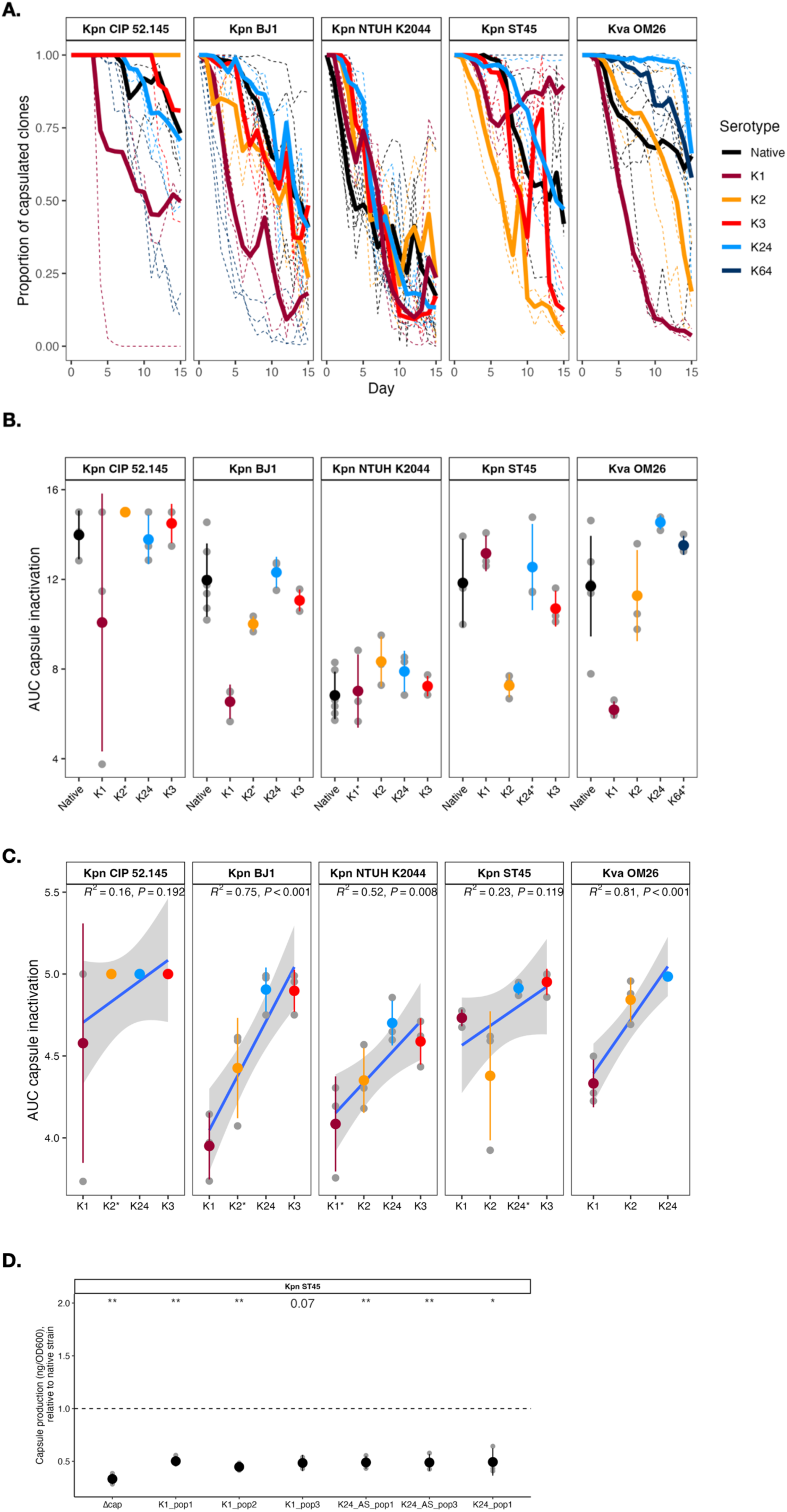
Experimental evolution of capsule-swapped strains. **A.** Proportion of capsulated clones through time after daily transfers in nutrient-rich medium. Independent replicates are indicated in dashed lines and bold lines correspond to the means of at least 3 independently-evolving populations. Evolution in nutrient-poor media is shown in Figure S6. **B.** Area under the curve (AUC) of the proportion of capsulated clones during the 15 days of the experiment in nutrient-rich medium. Grey dots correspond to independent populations and the colored dot indicates the mean. The native strain is shown in black. **C.** Linear regression (R^2^) between capsule serotype and area under the curve calculated from the first five days of the evolution experiment. **D.** Capsule production of intermediately capsulated clones, derived from different capsule-swapped ST45. Capsule production is portrayed as relative to their respective non-evolved (ancestral) strain. On the x-axis, the serotype is followed by the number of the population/replicate from which the clone was isolated from. Each black point represents the mean of at least three independent biological replicates. *p < 0.05; **p < 0.01; ***p < 0.001 one-sample t-test. Capsule quantification of the ST45 acapsular mutant (Δcap) is included as comparison. All capsulated clones have a mutation in Tat secretion system (Table 2). Similar observations were made in other genetic backgrounds and with other serotypes.

Towards the end of the experiment, capsulated clones increased in several populations of NTUH K2044, ST45, and BJ1 carrying either native or swapped serotypes, and reaching up to ∼50% in frequency (Figure 4A). These clones produced significantly less capsule than their ancestors (Figure 4D), suggesting a common compensatory response to the cost of capsule production. Genome sequencing of intermediate clones revealed frequent *wcaJ* mutations in ST45 and either plasmid loss or *rmpA* mutations in NTUH K2044, both known to reduce capsulation^24,47,49^. Unexpectedly, both strains also showed parallel mutations in the *tat* operon, including a nonsynonymous SNP in *tatC* (Table S3), an exporter of folded proteins, as well as in other genes (*zipA*, *envC*). These mutations, along with the absence of some TatC substrates in the periplasmic space, are expected to impair proper cell segmentation after cell division, leading to cell-chain formation.

Our data indicate that adaptation to an environment in which capsules are costly follows the same dynamics in native and swapped strains, suggesting that mitigating the generic cost of capsule production is larger than the cost imposed by a specific and novel serotype.

### K1 serotype determines virulence whereas resistance to biotic stress is dependent on the genetic background

One of the hallmarks of the plug-and-play model is that the newly inserted element should retain the same function across different genetic backgrounds. To address this, and given the importance of the capsule in *Kpn* epidemiology and pathogenicity, we tested different virulence-associated traits across the capsule-swapped strains. More specifically, we evaluated strain performance across representative life stages of *Kpn*: survival through the gastro-intestinal tract –including resistance to bile salts and oxidative stress, which can disrupt bacterial membrane^50^–, colonization ability, (i.e. biofilm formation), and hypermucoviscosity, which is associated to hypervirulence.

Insertion of any given serotype did not result in differences in resistance to physiological concentrations (0.05%) of primary (sodium cholate) and secondary (sodium deoxycholate) bile salts (Figure 5A and Figure S7A), or resistance to oxidative stress, assessed using 5 and 10mM of hydroxide peroxide (Figure 5B and Figure S7B-C).

**Figure 5.**
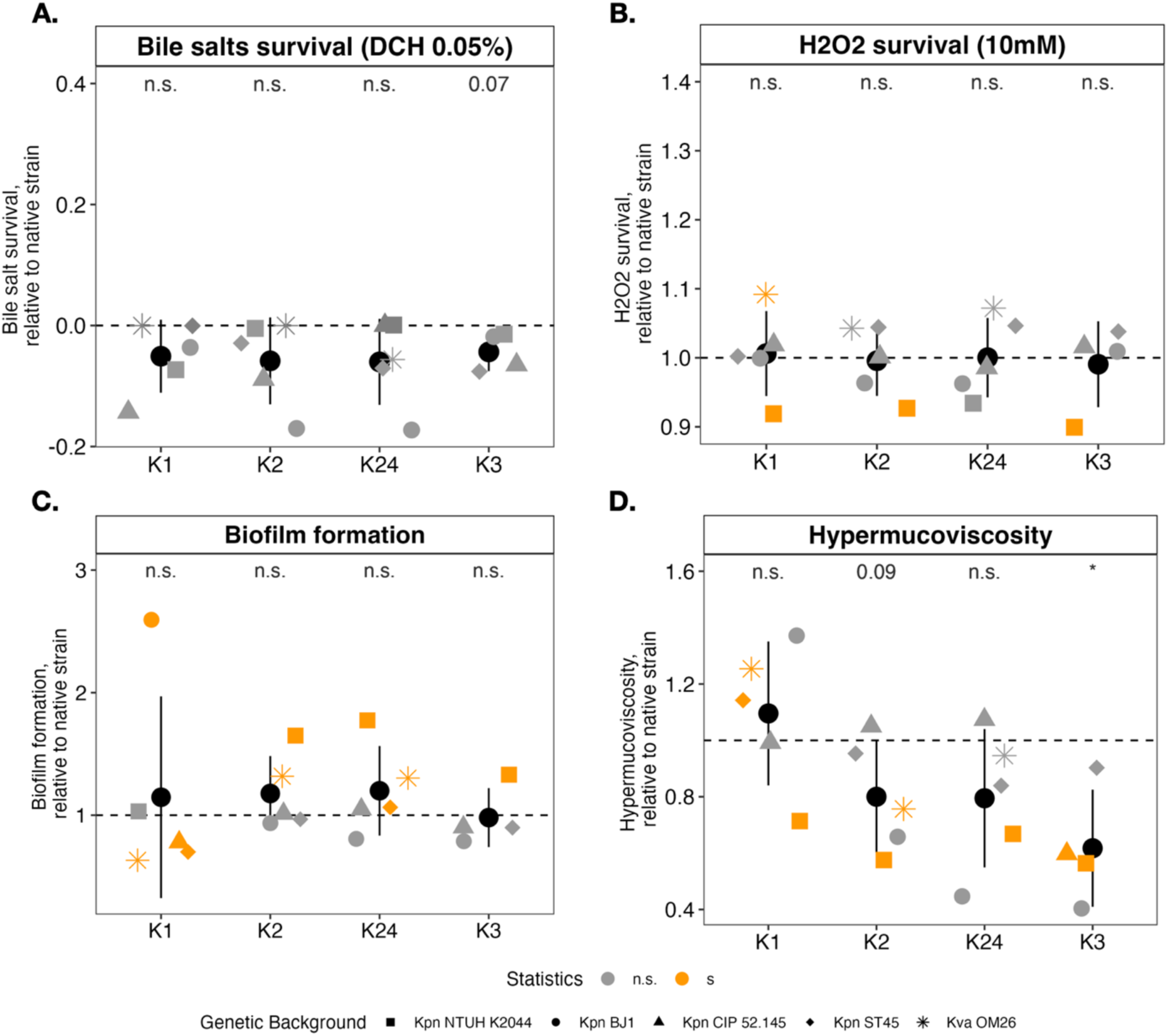
Virulence-associated traits of capsule-swapped strains. Survival of capsule-swapped strains to 0.05% deoxycholate (**A**) and to 10mM H2O2 (**B**), the ability to form biofilm (**C**), or hypermucoviscosity index (**D**) relative to their respective native strain. Each point represents the mean of at least three independent biological replicates of each capsule-swapped strain. The colour (orange or grey) indicates significant differences or not, relative to their own native strain. Shape of dots correspond to the different native strains (genetic background). The average across serotypes is indicated by the black dot (N=5). * < 0.05; **p < 0.01; ***p < 0.001, One-sample t-test, difference from 1.

Using a microtiter plate colorimetric assay, we tested biofilm formation in nutrient-rich (LB) and –poor (M02) media. Overall, biofilm is mostly driven by the genetic background, irrespective of the growth media (Figure S7D). Yet, in all strains but one, introduction of K1 serotype reduced biofilm formation. Conversely, the replacement of K1 by any other serotype increased biofilm formation (Figure 5C and Figure S6D). This suggests that thicker K1 capsules hinder adhesion potentially by masking fimbriae at the cell surface^51,52^.

A major capsule-associated phenotype is hypermucoviscosity (HMV). Typically linked to the presence of the *rmp* locus^49^, it has been mostly observed in K1/K2 serotypes. Recent work also associates HMV with other serotypes including K3^53^. Thus, we tested whether HMV is determined by the serotype, the strain or the presence of *rmp*. In the capsule-swapped-strains, three genetic backgrounds (NTUH K2044, BJ1 and CIP 52.145) are rmp+, while two backgrounds are rmp-(ST45 and OM26). Stepwise regression analysis identified serotype as the strongest predictor of HMV (Table S1 and S2). In rmp-strains*, Kpn* ST45 and *Kpn* OM26 (originally KL24 and KL64 respectively), introduction of the K1 capsule locus significantly increased HMV (Figure 5D). In *Kpn* BJ1 and *Kpn* CIP 52.145 –both K2 and rmp+– replacement of the K1 locus enhanced HMV but not significantly so. Conversely, replacement of the native K1 with any other serotype in *Kpn* NTUH K2044 reduced HMV (orange squares-Figure 5D). Finally, introduction of K3 serotype, not only did not increase HMV, but significantly reduced it.

Collectively, our results show that K1 capsules have conserved properties across different genomic backgrounds, resulting in limited biofilm formation and an increase of hypermucoviscosity, even in the absence of *rmp* (Figure 5C and Figure S8). This underscores how capsule swaps can directly shape *Kpn* interactions with the host and influence its in-patient evolutionary trajectory. It further shows modularity of the capsule as serotype-specific phenotypes are maintained upon transfer and integration in various genetic backgrounds.

## DISCUSSION

Our study investigates the evolution of complex functions and the consequences upon integration into a new host genome. The pervasive exchangeability of capsule serotypes within and across species is mostly driven by biotic stresses, host immunity, phage predation and nutrient availability^24,30,54^. Such exchangeability implies an underlying modularity of the genome architecture, allowing fast adaptation to changing conditions. Indeed, for capsule swaps to be advantageous, these exchanges should result in minimal fitness costs, whilst conserving their functionality (and thus, expression) across different genetic backgrounds. Here, we undertook an integrative approach and analyzed a collection of capsule-swapped strains to present evidence for a clear example of plug-and-play evolutionary dynamics in Bacteria.

First, despite the tight link between the capsule and the central carbon metabolism^44,45^, no regulatory network rewiring was observed. Transcriptomic analyses revealed minimal gene expression changes upon introduction of a novel serotype. The expression of the capsule locus itself does not change across the different genetic backgrounds but is mostly specific to each serotype (PCA-Figure S3). Accordingly, capsule production followed a conserved hierarchy across serotypes –K1 > K2 > K24 > K3–, regardless of the host genome. A similar conserved hierarchy existed across genetic backgrounds, regardless of serotypes. Indeed, the three genetic backgrounds associated with the highest capsule production, are hypervirulent strains (NTUH K2044, BJ1, and CIP 52.145). This suggests a very tight regulation of capsule genes, dictated both by the serotype and by the genetic background, but with limited, or no interaction (Table S3).

Second, both growth measurements across different environments and direct competitions show no significant differences between capsule-swapped strains compared to their native counterparts. These findings align with prior work showing that most HGT events have only minor fitness effects under laboratory conditions when integrated at a neutral position^55^. This is also in line with the evolutionary dynamics observed during the first steps of co-adaptation upon acquisition of a novel serotype. Specifically, our evolution experiment shows that the maintenance of a serotype depends more on environmental conditions (nutrient availability) than on a shared life-history between the capsule locus and its genetic background. Indeed, we show that all capsulated strains follow similar evolution trajectories suggesting that novel capsules are ready-to-use and require no co-adaptation to the genome. Additionally, the molecular mechanisms underlying adaptation to novel environments are similar in both capsule-swapped and their respective native strains, as shown by mutations emerging in parallel across all strains. These mutations result in the modulation of capsule expression, without altering the capsule biochemistry. Our data shows that capsule swaps have surprisingly little impact on the expression of other functions. Furthermore, the serotype specificities are maintained across genetic backgrounds. This modularity, relative to the rest of the genome, contributes to explain the high rates of swaps found in natural populations.

Finally, key phenotypes and functions associated to capsule serotypes should persist across genetic backgrounds. An early study in which capsule-swapped were generated by recombining large chromosomal segments encompassing the capsule locus^37^ showed that not all virulence-associated traits were inherited with the serotype and postulated that the role of capsule in *Kpn* virulence is likely multifactorial and dependent on epistatic interactions with the genetic background^37^. Our work, in line with Huang and colleagues, who also performed exchanges of the capsule locus in *Klebsiella*^34^, clearly shows that some capsule locus alone can encode specific functionalities. For example, a specific capsule locus of K1 alone, independent of the presence of the *rmp* locus, generically increases HMV of the strain. Similar conclusions were drawn when testing survival profiles to macrophages and in a liver infection context using capsule-swapped strains, where virulence levels –high or low– were dictated by serotype alone, irrespective of genetic background^34^. Among all other phenotypes tested, we could not observe any other specific serotype-function associations. Expanding the panel to include more ecologically diverse serotypes may uncover other serotype-specific functions. Also, more complex selection pressures encountered *in vivo,* such as competition within complex microbial communities or growth under anaerobic conditions, may reveal novel serotype-specific functions. Indeed, we had previously shown that exchange of serotype could also result in exchange of susceptibility against phage infection^39^. Hence, capsule swaps very specifically change some bacterial phenotypes without affecting other processes.

In our study, we did however detect few and limited context-dependent epistasis. Introduction of the same K1 capsule locus into two K2 genetic backgrounds (BJ1 and CIP 52.145) resulted in contrasting outcomes: minimal transcriptomic changes in *Kpn* BJ1 versus widespread regulatory shifts in *Kpn* CIP 52.145. More striking, despite numerous attempts, we were unable to introduce the K3 serotype cloned from *Kpn* ATCC13883T into *Kva* OM26 strain suggesting that some capsule swaps may be highly deleterious or even lethal. Such incompatibility may reflect underlying differences in core metabolic gene content, as each *Klebsiella* taxon harbors a distinct metabolic profile^56^. Indeed, both K2 strains as well as *Kpn* ATCC13883T and *Kva* OM26 belong to different phylogenetic sublineages and subspecies, respectively. Each species and sublineage has a distinct metabolic profile^56^, suggesting that capsule-host compatibility may depend on specific metabolic pathways that help mitigate negative epistasis. However, both examples of epistasis may be a consequence of the experimental set up. Capsule-swapped strains were constructed through allelic replacement by recombination at conserved flanking regions (*galF* and *ugd*), yet, in natural populations, we observed that recombination tracts often extend well beyond the capsule locus^24^. These extended tracts may result in co-transfer of additional loci, such as the O-antigen biosynthesis enzymes encoded on the *rfb* locus^57^ just downstream the capsule locus, the *his* locus^58^ or other core genes. Co-transfer would ensure direct surface structure compatibility, membrane homeostasis, and metabolic integration. A recent study showed that capsule production can be heterogeneous within clonal populations, a variability shaped by both the capsule locus and the genetic background (Nucci et al. 2025). While insertion of the K1 locus consistently resulted in non-heterogeneous phenotype across genetic background, other serotypes did not follow a consistent pattern, indicating that some capsule-encoded traits might be governed by epistasis (Nucci et al. 2025). Nevertheless, examples supporting this model –where capsule integration is restricted by host-genome compatibility– remain rare cases.

Our findings support a plug-and-play model of capsule evolution, in which capsule loci act as modular transferable elements. This contrasts with capsule evolution in *Streptococcus pneumoniae* (*Spn*), where fitness outcomes of capsule-switched isogenic strains are often shaped by strong serotype-genotype epistasis^59^. These differences could be due to contrasting evolutionary mechanisms for capsule variation. In *Spn*, capsules typically change by intra-locus recombination, generating mosaic or chimeric loci rather than full-locus novelty ^59^. Although some modularity exists—for instance, flippases show partial interchangeability—this is constrained by substrate specificity^60^. Relaxed specificity can lead to growth defects after recombination. These are likely counter-selected in nature^61^, limiting the spread and persistence of deleterious chimeras. Finally, the narrow ecological niche of *Spn* may result in much stronger selection than the more environmentally versatile *Kpn*. Hence, the mechanism of capsule change in *Spn*, intralocus recombination, appears to impose greater evolutionary constraints than the whole-locus replacement observed in *Kpn*.

Taken together, the study of a complete macromolecular system rather than isolated gene interactions allowed us to provide an explanation for the pervasive transferability of capsule loci not only across distant *Klebsiella* lineages but also between species and genera. Collectively, our data show that capsule loci can be horizontally transferred and successfully integrated into diverse genetic backgrounds with minimal disruption to cellular fitness or regulatory networks. Further, capsule loci act as modular elements with inherent functional properties which are inherited across genetic backgrounds. Their large diversity and rapid evolution do not seem constrained by epistasis, but instead integrate and persist across diverse genetic backgrounds, exemplifying a true plug-and-play model of evolution.

## MATERIALS & METHODS

### Bacterial strains and growth conditions

*Klebsiella spp*. strains were grown at 37°C in 4 mL Luria Bertani Broth (LB Miller) – unless indicated otherwise – in 14 mL tubes and under orbital shaking conditions (250 rpm) or agar plates. Nutrient-poor medium corresponds to minimal medium M63B1 supplemented with 0.2% of glucose (M02). The strains used in this study, as well as their genomic annotations and references, are described in Supplementary Data S1.

### Scarless serotype swap

The scarless serotype swaps was performed as described before ^39^. ***Capsule deletion mutant.*** Capsule deletion was performed by recombination at the homologous regions (5’-*galF* and *ugd*-3’). The different genetic backgrounds, referred to as native strains in the following work, were transformed by electroporation with a λ-red-carrying plasmid (pKOBEG199, see Supplementary Data S2, *Plasmids*). Transformants were selected on LB plates supplemented with tetracycline and 0.2% of glucose. Competent cells of transformants were induced with 0.2% L-arabinose for 2 hours, to increase recombination upon transformation by electroporation of the deletion cassette leading to loss of the capsule locus and replacement with the deletion cassette –containing a kanamycin resistance gene, one I-Scel cut site and two FRT sites– and selected on LB with kanamycin at 37°C. Non-capsulated colonies were then selected and transformed by electroporation with pMPIII, a plasmid encoding a flippase (FLP) (see Supplementary Data S2, *Plasmids*). The kanamycin resistance gene was excised by the FLP flippase acting on the FRT sites leaving a small scar containing a I-Scel cute site for the swap step. Cells were plated on LB supplemented with spectinomycin and incubated at 30°C. A colony was then grown overnight at 42°C to cure the pMPIII plasmid then plated on LB. Non-capsulated clones that grow only on LB were selected. ***Generating pKAPTURE vectors***. Capsule cloning was done using a linear cassette named pKAPTURE consisting of two homolog regions in reverse (5’-*galF* and *ugd*-3’), an origin of replication, a kanamycin resistance gene and two I-Scel cut site. The cassette circularizes around the capsule locus *via* recombination and capture the whole locus to form a circularized pKAPTURE. To do so, the linear cassette was transformed in electrocompetent pKOBEG199-strain, in which expression of l-red had been induced (see above). Cells were then plated on LB with kanamycin and incubated at 37°C. Kanamycin-resistant transformants are expected to carry recircularized pKAPTURE containing the capsule of the native strain. These transformants were then grown overnight supplemented with kanamycin and EDTA. pKAPTURE plasmids were extracted and electroporated into the abovementioned capsule deletion mutant to generate a stock of pKAPTURE expressing a given capsule serotype. After recovery, cells were plated on LB supplemented with kanamycin and allowed to grow overnight at 37°C. Capsulated colonies identified were restreaked in parallel on LB and LB supplemented with kanamycin (50 μg/mL). Capsulated colonies on LB supplemented with kanamycin but non-capsulated on LB were selected and considered a source of pKAPTURE vector which carries the capsule locus of interest. ***Generating capsule-swapped strains.*** Electrocompetent capsule deletion mutants were transformed with pKAPTURE encoding a specific capsule locus. After recovery, cells were plated on LB with kanamycin and grown ON at 37°C. To allow integration of pKAPTURE, electrocompetent cells of the strain with the pKAPTURE were transformed with pTKRED (plasmid encoding RecA and I-Scel enzymes, see Supplementary Data S2, *Plasmids*). Cells were recovered at 30°C in LB supplemented with kanamycin, to avoid pKAPTURE loss, and 0.2% glucose for 1h30, prior to plating on LB with kanamycin, spectinomycin and 0.2% glucose and incubated at 30°C. To induce integration, individual colonies were resuspended in M63B1 supplemented with spectinomycin, 0.2% L-arabinose, 0.2% glycerol and grown at 30°C. I-Scel endonuclease cuts the chromosome and the circularized pKAPTURE containing the capsule locus at the I -Scel sites and *recA* expression increased recombination leading to chromosomal repair by of the capsule locus carried by the pKAPTURE. After 12 to 24 hours, cells were diluted depending on culture turbidity, plated on LB supplemented with 0.2% glucose and grown at 42°C to cure pTKRED. Capsulated colonies identified were restreaked in parallel on i) LB, ii) LB with kanamycin, and iii) LB with spectinomycin. Capsulated clones that only grow on LB are considered successful swapped clones. Swapped clones were finally verified by Illumina sequencing. Complementation strains have been performed as controls and will be referred as complemented strains.

### Capsule extraction and quantification

The bacterial capsule was extracted as described before ^62^ and quantified by the uronic acid method^63^. Briefly, OD_600_ of overnight cultures in LB medium was measured. Then, 500 μL were transferred to an Eppendorf tube with Zwittergent and were centrifuged. The supernatant was discarded as to specifically measure cell-bound surface polysaccharides, and washed with ethanol, and dissolved in double-distilled water. The uronic acid concentration of each sample was determined from a standard curve of glucuronic acid. Finally, to normalize for number of cells, the concentration was then divided by the OD600nm of the overnight culture.

### Growth curves

Overnight cultures were diluted at 1:100 in the different growth environments. 200μL of each subculture was transferred in a 96-well microtiter plate and allowed to grow at 37°C, under orbital shaking for 16 hours. Absorbance (OD_600_) of cell cultures was measured every 15 minutes with a TECAN Genios^TM^ plate reader.

### Hypermucoviscosity

Precultures of each strain were grown overday in LB and then diluted to 1:200 in M02 for overnight culture. To initiate the experiment, cultures were vigorously vortexed and 200μL of culture were transferred into a 96-well microtiter plate. Absorbance (OD_600_) was measured to set the initial OD (OD_i_). Cultures were then sedimented by slow centrifugation (2500 rpm) for 5 minutes at room temperature. 200μL of the top part of culture were then transferred into a 96-well microtiter plate. Absorbance (OD_590_) was measured to set the final OD (OD_f_). The hypermucoviscosity index was calculated as (OD_f_/OD_i_).

### Biofilm formation staining using crystal violet

Strains were grown overnight in LB and diluted at 1:100 in the different growth environments. 200μL of each subculture was transferred in a 96-well microtiter plate then incubated at 37°C for 24 hours. Supernatant was removed by flicking the plate and bacteria were washed thrice with water. Extra water was emptied by flicking. The surface-attached bacterial mass remaining in the well was stained with 220μL of crystal violet (1%) for 30min at room temperature, washed thrice with water and air-dried. The bacterial mass was resuspended in 220μL of ethanol:acetone (80:20) and absorbance (OD_590_) was measured.

### Bile salts survival assays

Overnight cultures in LB were serially diluted and plated on LB or on LB supplemented with either 0.05% (physiological conditions) or 0.5% of CHO (primary bile salts, sodium cholate) or DCH (secondary bile salts, sodium deoxycholate). Colonies were allowed to grow at 37°C and surviving CFU were counted after 24 hours.

### Hydroxide peroxide survival assay

Overnight cultures in LB were diluted at 1:100 in LB and grown at 37°C until OD∼0.6-0.8. 500μL of overday culture were transferred in a 96-deep well plate supplemented with either 0, 5, 10 or 15mM of H_2_O_2_. Cultures were incubated for 1h at 37°C without shaking. Subcultures were serially diluted, spotted on LB agar plate and grown overnight at 37°C. Surviving CFU were counted after 24 hours and compared to the control without H_2_O_2_.

### Evolution experiment

Three independent clones of each strain were used to initiate each of the three evolving populations in nutrient-rich (LB) and nutrient-poor (M02) media. Each day, populations were diluted 1:100 into fresh media and allowed to grow for 24 hours at 37°C. In parallel, cultures were serially diluted, plated and visually inspected each day on LB to count capsulated and acapsulated clones. The experiment was performed for 15 days accounting for an estimated 100 generations (= 15 days x 6.7 generations/day) (based on *E. coli* generations/day). Although each growth media has slightly different carrying capacities, all cultures reached bacterial saturation before daily passaging, ensuring that the different populations underwent a similar number of generations across growth media.

### Fluorescent strains construction

Scarless integration of the *mCherry* fluorescent protein was performed by double recombination event at the neutral site attTn7 downstream of the *glmS* locus^64^. Briefly, upstream and downstream 500 bp regions of *glmS* were amplified by PCR, as well as the *mCherry* gene preceded by the strong constitutive pLpp promoter (See Supplementary Data S3, *Primers*). To assemble the fragments into a vector (pKNG101), we used GeneArt™ Gibson Assembly HiFi kit (Invitrogen) and incubated the mix for 30 minutes at 50°C. The assembly reaction was diluted 1:4 and electroporated into competent *E. coli* DH5α strain and selected on LB plates with streptomycin (100 μg/mL). Colonies with integrated fragments were checked by PCR, extracted, electroporated into *E. coli* MFD λ-pir strain, and used as a donor strain for conjugation in capsule-swapped strains. Single cross-over mutants (transconjugants) were selected on streptomycin plates (200 μg/mL) and double cross-over mutants were selected on LB without salt and supplemented with 5% sucrose after growth at room temperature. Mutants were verified for their sensitivity to streptomycin and by PCR, and *mCherry* expression confirmed by fluorescent microscopy.

### Bacterial competitions

Precultures of each strain were grown overday in LB and then diluted to 1:100 in M02. Strains with or without fluorescent tags were mixed in a 1:1 ratio. The mixes were diluted to the 1:100 either into M02 or cold PBS 1X to avoid further growth and verify cell ratio in the competition mix using flow cytometry (T_0_). The M02 plate was placed at 37°C in shaking conditions in a microtiter plate reader to allow bacterial competition. After 24 hours of growth (T_24_), samples were diluted into cold PBS 1X to the 1:100, and the proportion of each strain was assessed by fluorescence-activated cell sorting (FACS) analysis using CytoFlex S. Four competitions between each genotype were performed, two in which one of the strains was tagged *mCherry*, and the other two with the *mCherry* integrated in the other competitor.

### FlowJo analyses and fitness calculation

Flow cytometry data were analysed using FlowJo software (v10.10.0). Subsequent analyses were performed using R.4.4.1. The relative fitness was calculated by dividing the ratio of cells at T_24_ compared to T_0_. Competitions with a relative fitness <0.5 and >2, most likely indicative of a technical mistake, were removed. These accounted for 35 out of 346 competitions, evenly distributed across competitions. Control experiments to assess whether *mCherry* integration was associated to a cost revealed no difference in fitness (Figure S4B).

### RNA extraction and sequencing

***Extraction***. An overday culture was started in M02 from freshly plated colonies and incubated at 37°C until OD_600_ reached ∼0.6-0.8. RNA was extracted using the Macherey Nagel Trizol (ref:740971.250) according to the manufacturer’s instructions and treated with DNase I provided in the kit. RNA concentration, quality, and integrity from four independent replicates were checked using the Invitrogen Qubit and the Agilent 2100 Bioanalyzer system. Four independent samples per population were sequenced. ***Sequencing***. Library preparation and sequencing were carried out at the Biomics Platform, Institut Pasteur. cDNA libraries were prepared from 1–3 µg of total RNA using the Illumina Total RNA Library Preparation Kit (Illumina, USA), following the manufacturer’s protocol. To facilitate rRNA depletion, Ribo-Zero Plus Microbiome probes (Illumina) were used. Index barcodes were added by PCR for 13 cycles. Unbound adaptors and index primers were re-moved via purification with AMPure XP magnetic beads (Beckman Coulter, USA). The final libraries displayed an electrophoretic size distribution ranging from 250 to 900 bp, with a predominant peak at approximately 400 bp, as assessed on a 3500 Fragment Analyzer (Agilent Technologies, USA). Sequencing was performed on a NextSeq 2000 system using a P3 50-cycle flow cell (Illumina) to generate 67-nt single-end, dual-indexed reads.

### RNA sequencing analysis

***Cleaning.*** Single-end strand-specific 65 bp reads were cleaned of adapter sequences and low-quality sequences using cutadapt version 4.9^65^ with options “-m 25 -q 30 -O 6 --trim-n --max-n 1”. Gene expression quantification was performed using salmon version 1.9.0 with the “-l A” option^66^. For each condition, a specific reference transcriptome was built concatenating both the fasta transcriptomes of the genetic background and the capsular region. Seven out of 96 samples were excluded due to poor sequencing quality, and all strains were represented by at least three biological replicates. ***Gene expression analysis.*** Gene expression data were analyzed using R version 4.3.2^67^ and the Bioconductor package DESeq2 version 1.42.1^68^. The data structure was explored using a Principal Component Analysis based on the replicate-adjusted variance-stabilized transformed count matrix. Replicate adjustment was performed using the removeBatchEffect() function from the limma R package^69^. The normalization and dispersion estimation were performed using the default parameters and statistical tests for differential expression were performed applying the independent filtering algorithm. Four background-specific independent analyses were conducted to compare the gene expression across conditions (i.e. capsules). For each background strain, a generalized linear model, including the replicate effect as blocking factor, was set to test for the capsular swap effect on gene expression. For each pairwise comparison, shrinkage of the log_2_(FC) was performed using the ashr method^70^, raw p-values were adjusted for multiple testing according to the Benjamini and Hochberg procedure^71^ and genes with an adjusted p-value lower than 0.05 were considered differentially expressed.

Several quality controls were performed. First, Principal Component Analyses (PCA) on the expression of core capsule genes revealed that the first component could differentiate acapsulated strains from all other capsulated strains with high inertia (Figure S4B). Secondly, in Δcps strains, all capsule genes were found highly downregulated due to their absence. Thirdly, capsule genes which are specific to each of the native capsule locus and absent in the swapped strains were significantly downregulated. Conversely, upregulated capsule genes were mostly genes specific of the newly acquired capsule locus and not found in the native one. Finally, PCA using only the core capsule genes allowed the clustering of independent biological replicates together Figure S4D.

### Whole genome sequencing

The genomes of capsule-swapped strains and clones producing reduced capsule amounts were extracted using the guanidium thiocyanate method^72^ prior to Illumina sequencing. Their sequences were compared with ancestral genotypes using *breseq* v.0.35.7^73^ with default parameters. Some mutations *(tatC, wcaJ*) were further verified by PCR (See Supplementary Data S3, *Primers*) and subsequent Sanger sequencing. Full list of identified mutations is provided in Supplementary Data S4.

### Other software and package

All the data analyses were performed with R version 4.4 and Rstudio v2022.02.1, except when precised otherwise in Methods. For data frame manipulations, we used dplyr v1.1.4 along with the tidyverse packages v2.0.0. We used the packages ggpmisc v.0.6.0 and Kendall v2.2.1 for the linear regressions and fitness transitivity, respectively. Graphs were performed using ggplot2 v.3.5.1, gridExtra v.2.3 and ggtext v.0.1.2.

## ACKNOWLEDGEMENTS

We are grateful to Amandine Nucci for technical aide with RNA extractions. We also thank Y. Vitrenko, L. Lemée and E. Kornobis of the Biomics Platform, C2RT, Institut Pasteur, Paris, France, supported by France Génomique (ANR-10-INBS-09) and IBISA for the samples QC, libraries and sequencing. We thank P.H. Commère, S. Schmutz, S. Novault of the Institut Pasteur Flow Cytometry Platform for the training, help and guidance. We thank Fabienne Benz for the gift of the *mCherry* gene, and Jean-Marc Ghigo and Christophe Beloin for providing lab space and support.

We thank Benoit Pons, Matthieu Haudiquet and Samay Pande for critical reading of the manuscript and Basile Beaud Benyahia and Cyril Anjou for scientific discussion.

The artificial intelligence ChatGPT was used to improve writing style and grammar of the manuscript.

---

**Figure S1.**
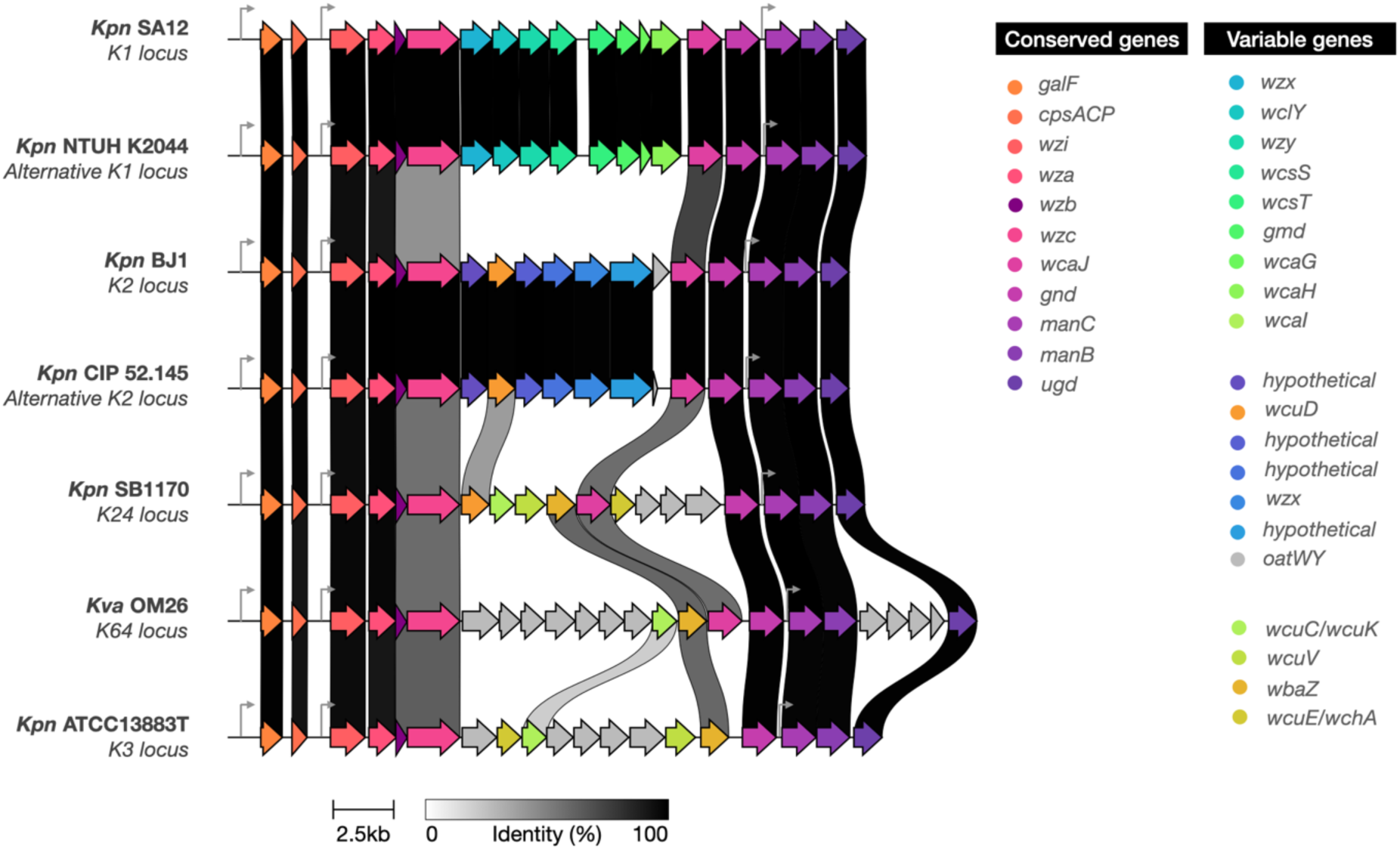
Genomic organization of capsule loci of strains used in the study. Capsule locus identification and annotation were done using Kaptive (Stanton et al. 2025). Alignment and visualization were done using Clinker (Gilchrist and Chooi 2021). Grey arrows indicate promoters of the locus. Alternative loci (from *Kpn* NTUH K2044 and *Kpn* CIP 52.145) are native loci but not used as template to generate the capsule-swapped strains. Figure was generated using clinker (https://github.com/gamcil/clinker) with modifications.

**Figure S2.**
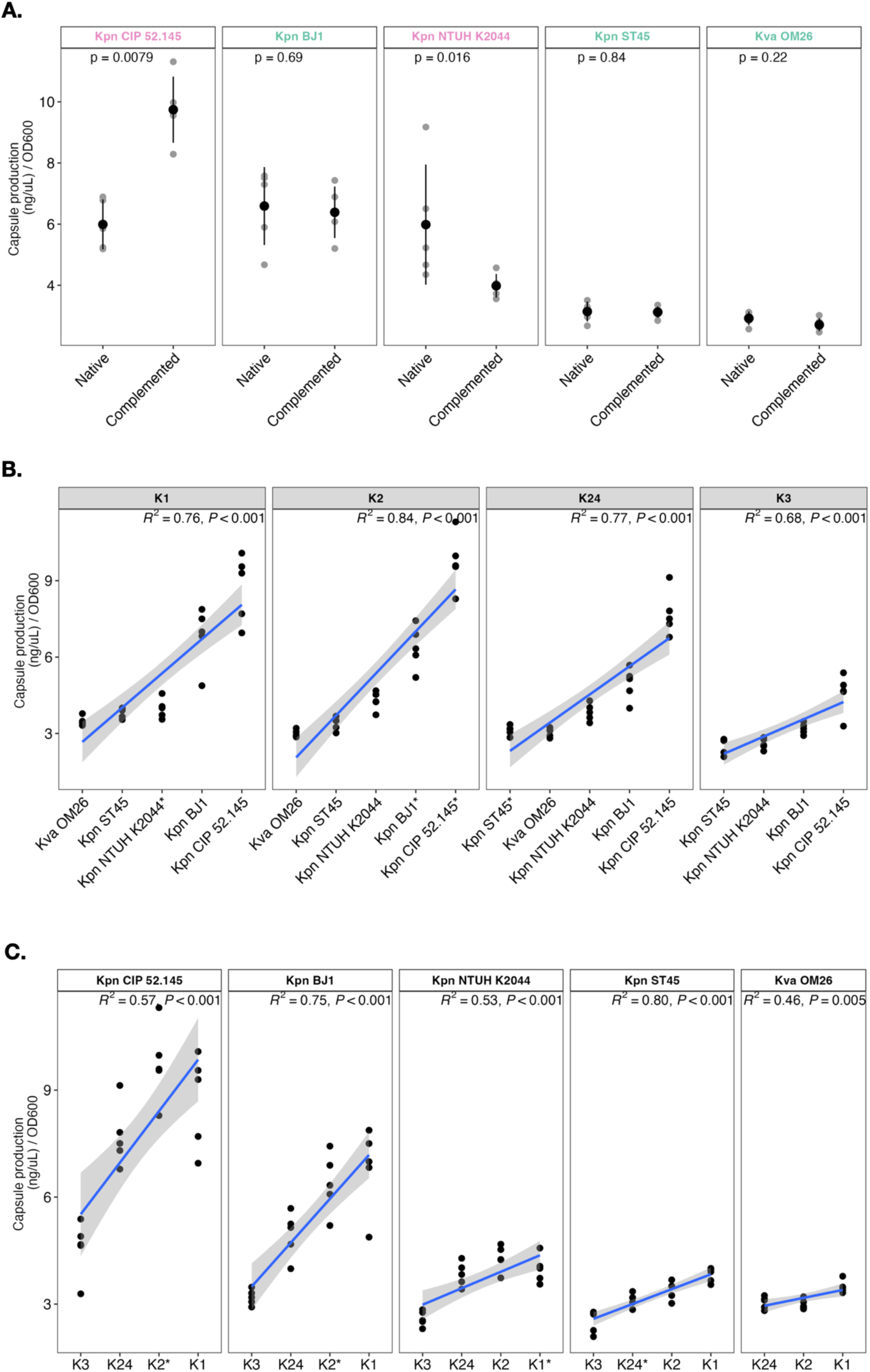
Capsule production of capsule-swapped strains determined by the uronic acid method. **A.** Comparison of capsule production between the wild type strain and its complemented strain corresponding to the reintroduction of either the native capsule locus (green) or an alternative capsule locus of the same serotype (pink). Values were normalized by an OD_600_ of 1. P-values correspond to unpaired Wilcoxon test. **B and C.** Capsule production across different capsule loci (**B**) or genetic background (**C**), ranked from lowest to highest capsule production. Blue lines represent regressions for each serotype (R^2^), and the surrounding grey area indicate the standard error. Each dot corresponds to an independent biological replicate.

**Figure S3.**
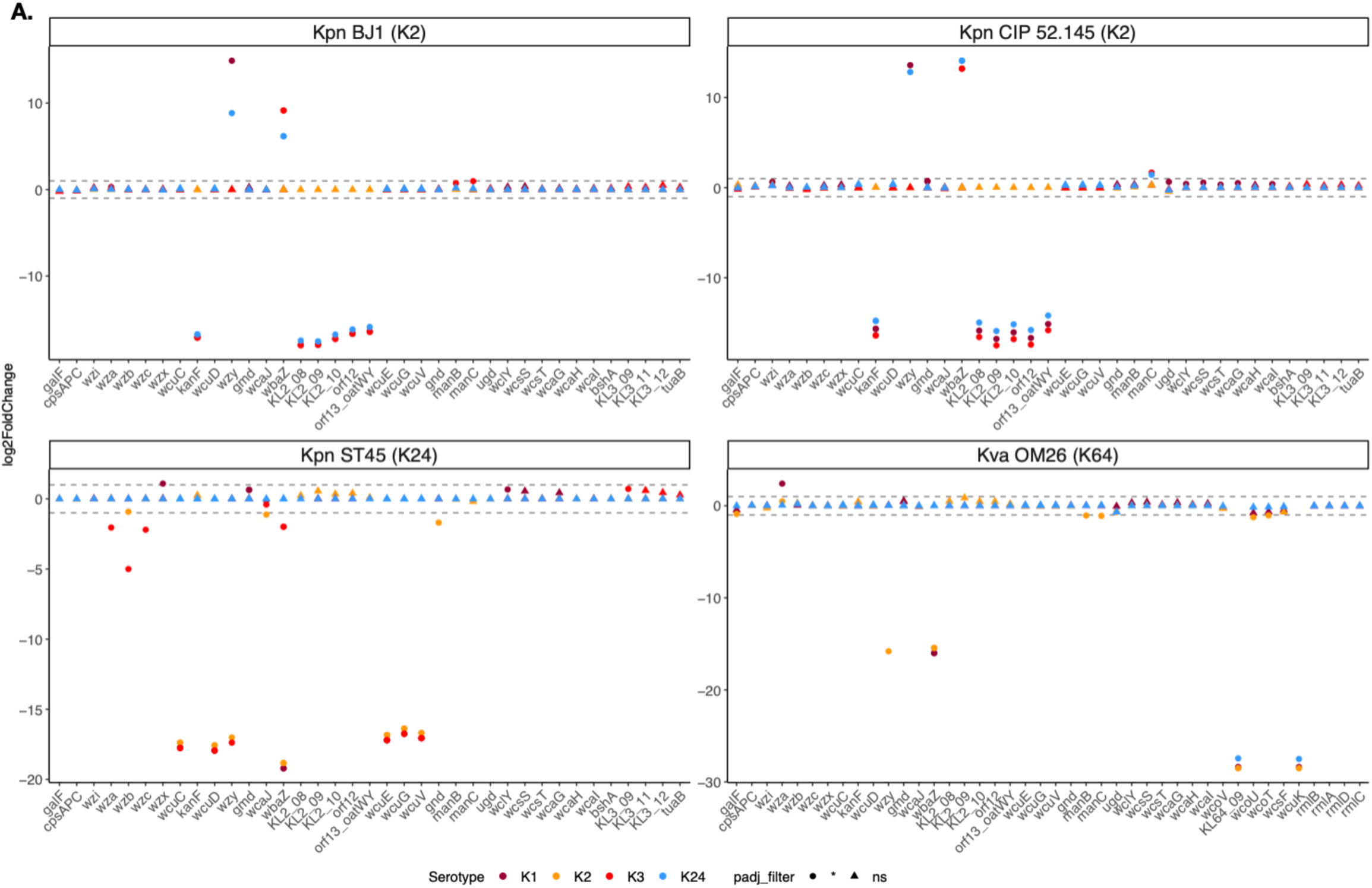
Transcriptomic analyses of capsule loci genes in capsule-swapped strains. Log_2_FC (fold change) of capsule locus genes, i.e. genes present in any of the five capsule loci type considered. The shape indicates the significance of the adjusted p-value and the color represents the capsule serotype. The genetic background and the native capsule serotype are indicated at the top of each subpanel.

**Figure S4.**
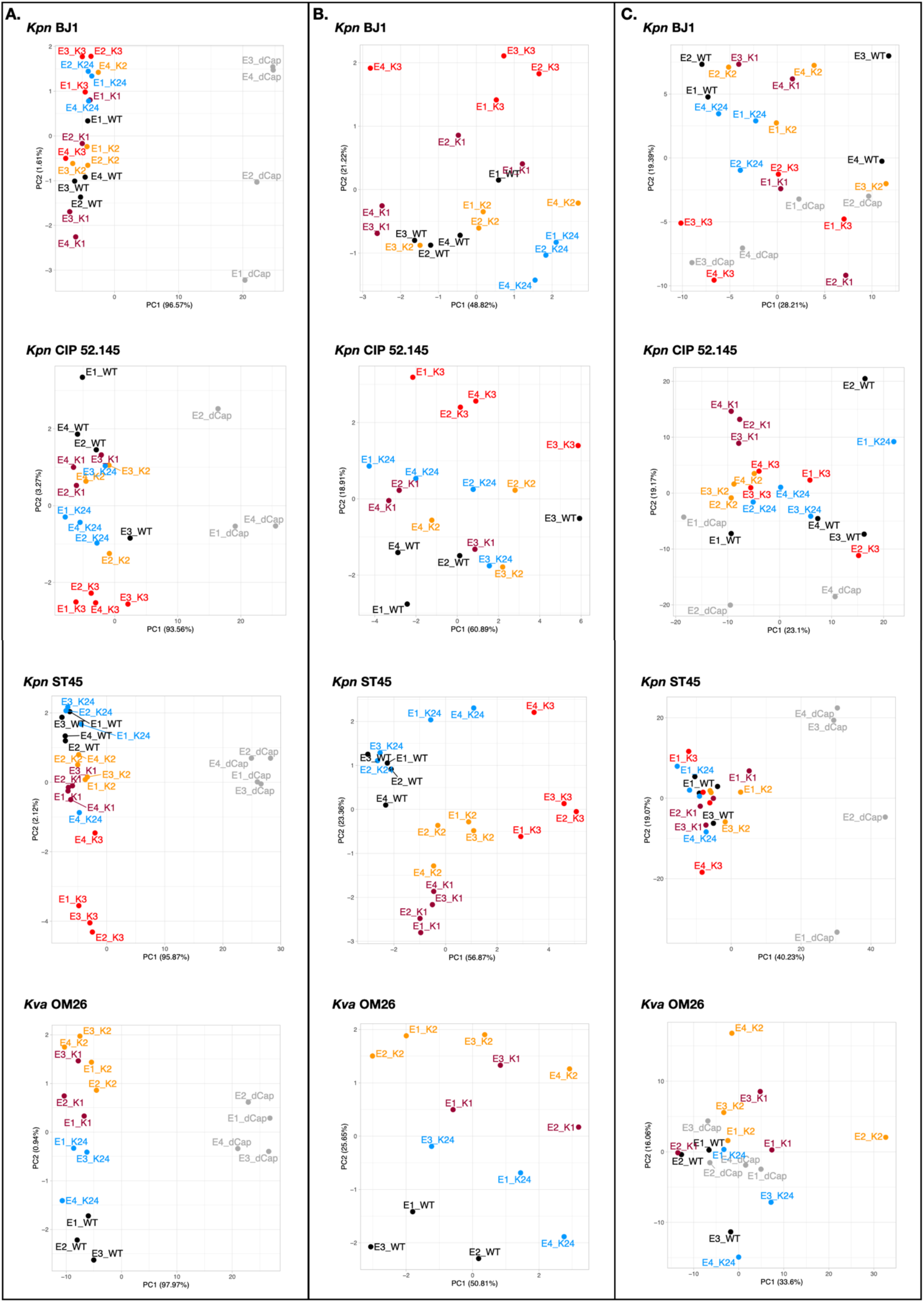
Adjusted Principal Component Analysis of gene expression. Components 1 and 2 are displayed and are followed by the percentage of variance explained by each component. Each dot represents a biological replicate annotated as follows: replicate number (EX), underscore and serotype (KX). PCA analysis was performed on the core capsule genes (**A-B**) and the core genome (**C**). For better visualization of the clustering of capsule-swapped strains, the PCA on the core capsule was done with (**A**) and without (**B**) taking into account the dCap (non-capsulated) strains.

**Figure S5.**
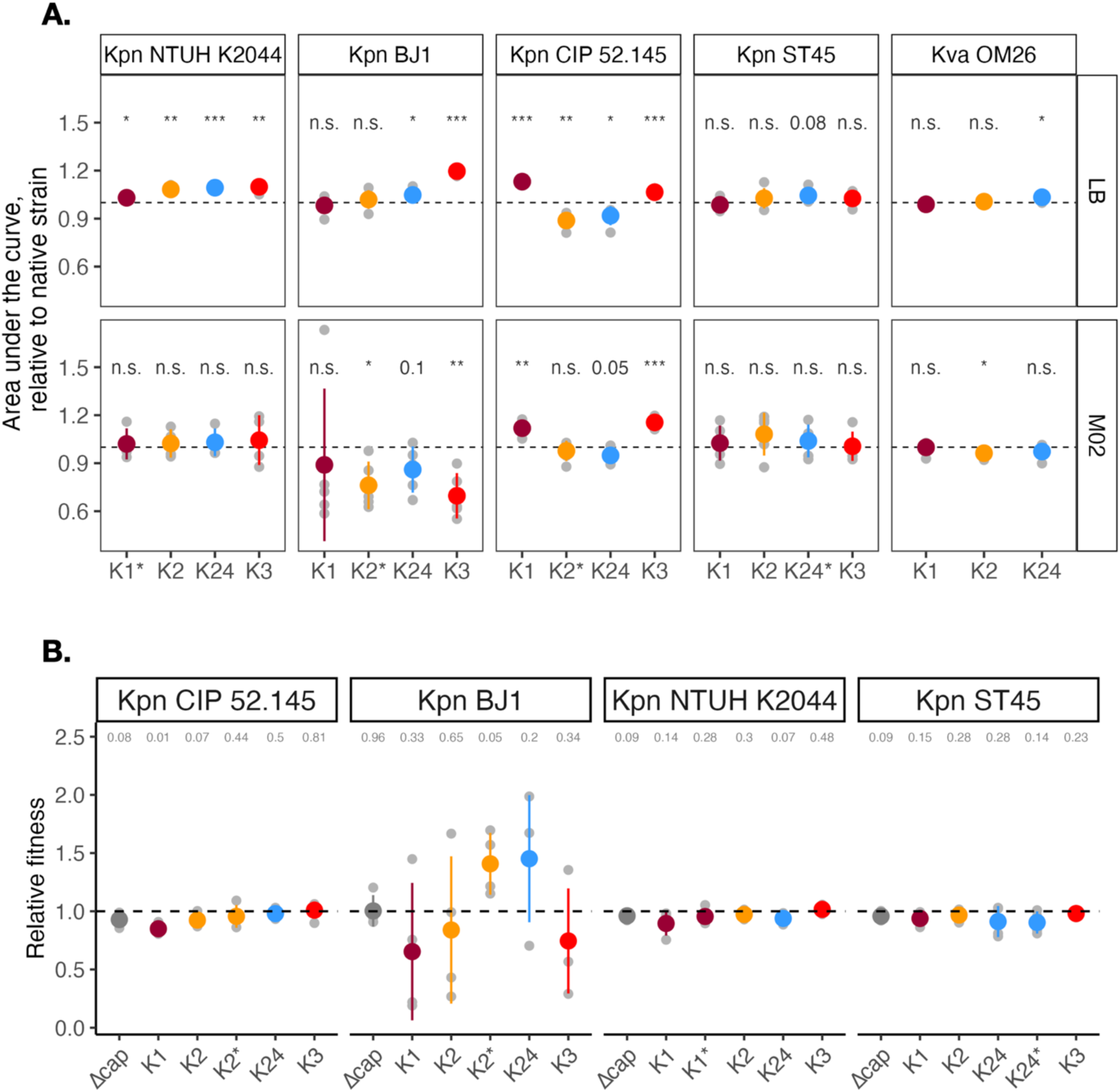
Growth and fitness of capsule-swapped strains. **A.** Area under the growth curve (AUC) of capsule-swapped strains relative to their respective native strain (dotted line) in either nutrient-rich (LB) or nutrient-poor (M02) media. Grey dots represent individual biological replicates (N=5). **B.** Pairwise competitions in nutrient-poor medium (M02) of fluorescent versus non-fluorescent strain (one-sample t-test, difference from 1). Asterisk beside K loci (*) indicates the native serotype of each strain. Δcap indicate non-capsulated control strains.

**Figure S6.**
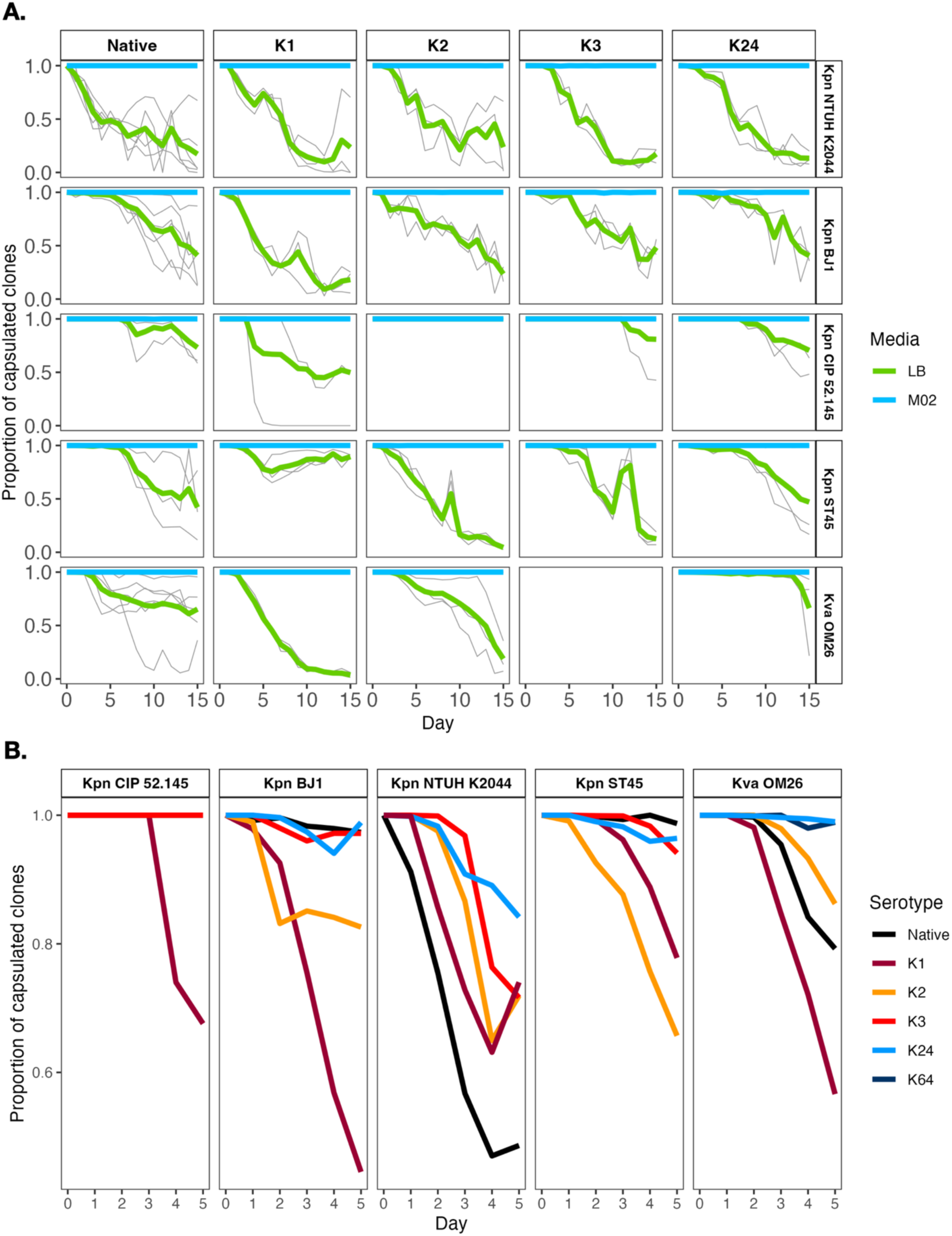
Evolutionary fate of capsule in the capsule-swapped and native strains. **A.** Proportion of capsulated clones throughout the fifteen days of evolution of parental strains and their respective isogenic capsule-swapped strains before daily passages of each culture either in nutrient-rich (LB, green line) or nutrient-poor (M02, blue line) media. Bold lines represent average of the independent populations of the same strain and environment. Grey lines represent each of the independent populations. **B.** Proportion of capsulated clones throughout the first five days of evolution in nutrient-rich medium. Each line represents the average of at least three independently evolving populations.

**Figure S7.**
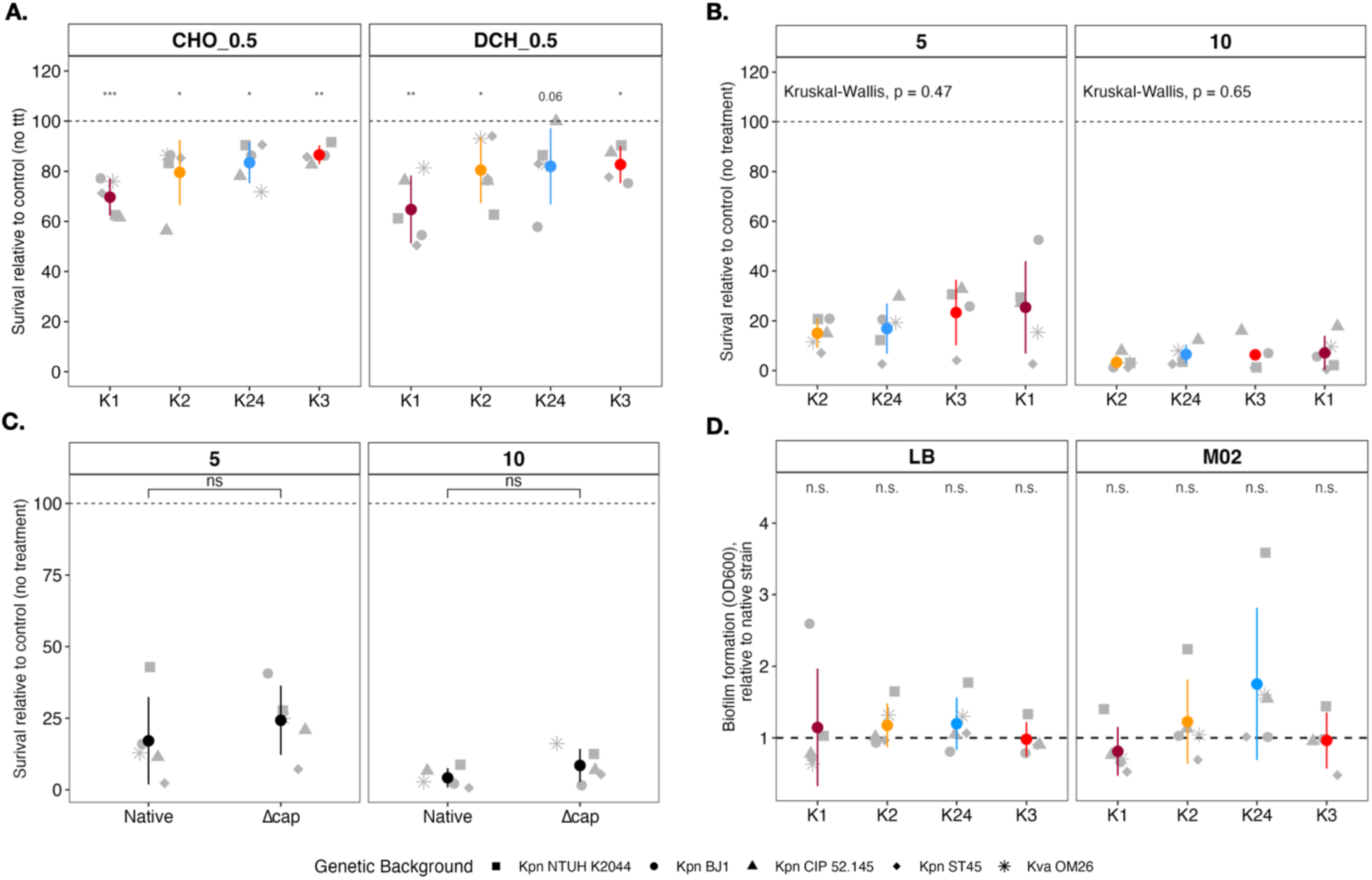
Survival to biotic stress of capsule-swapped strains. **A-B.** Capsule-swapped strains survival to 0.5% cholate (CHO) or deoxycholate (DCO)(**A**) and to 5mM or 10mM H_2_O_2_ (**B**), relative to their respective untreated condition. *p < 0.05; **p < 0.01; ***p < 0.001, one-sample t-test, difference from 100. **C.** Native capsulated strains and their respective acapsular mutant (Δcap) survival to 5 or 10mM H_2_O_2_, relative to their respective non-treated condition. ns, non-significant, two-sample paired t-test. **D.** Biofilm formation of capsule-swapped strains in nutrient-rich (LB) or nutrient-poor (M02) media, relative to their respective native strain. Shape of dots correspond to the genetic background; the serotype is indicated on the x-axis and by the color. Each point represents the mean of at least three independent biological replicates. ns:non-significant; one-sample t-test, difference from 1.

**Figure S8.**
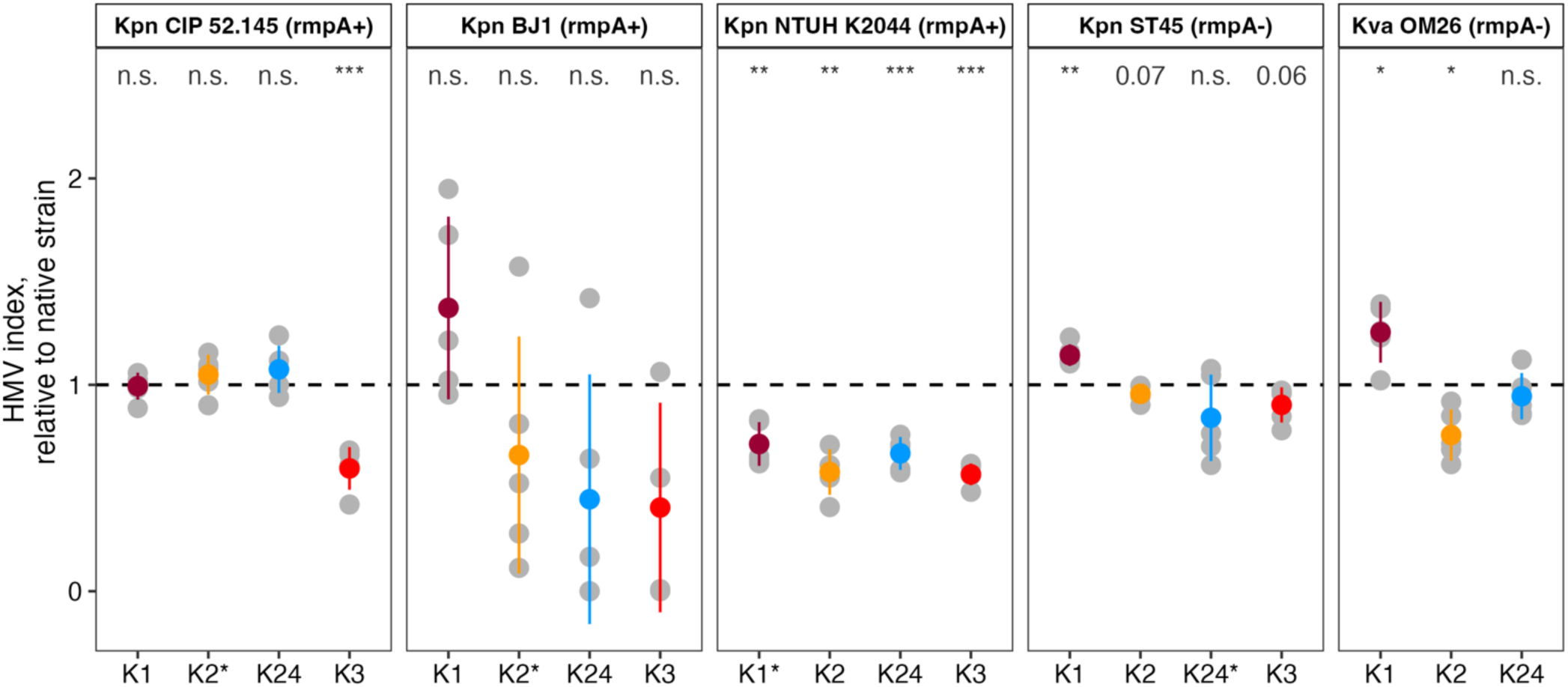
Capsule-swapped strains hypermucoviscosity index. Capsule-swapped strains hypermucovsiscosity index relative to their respective native strain in nutrient-poor medium. The serotype is indicated on the x-axis and by the color. Grey points represent independent biological replicates and color points represent the mean of these biological replicates. Asterisk beside K loci (*) indicates the native serotype of each strain. ns:non-significant; one-sample t-test, difference from 1.

**Table S1.** Statistical analyses – Multifactorial ANOVA for all traits analysed.

| **Trait** | **Variable** | **F value** | **P value** |
| --- | --- | --- | --- |
| *Capsule production* | Serotype | 55.39 | < 2e-16 *** |
|  | Genetic Background | 196.49 | < 2e-16 *** |
|  | Serotype::Background | 7.89 | 5.92e-09 *** |
| *Growth* | Serotype | 2.092 | 0.104 |
|  | Genetic Background | 7.114 | 2.96e-05 *** |
|  | Serotype::Background | 2.552 | 5.56e-03 ** |
|  | Media | 12.261 | 6.16e-04 *** |
|  | Background::Media | 12.532 | 8.95e-09 *** |
| *Growth in LB* | Serotype | 24.62 | 4.51e-11 *** |
|  | Genetic Background | 12.21 | 1.24e-07 *** |
|  | Serotype::Background | 17.14 | 5.14e-16 *** |
| *Growth in M02* | Serotype | 0.575 | 0.633 |
|  | Genetic Background | 9.668 | 2.58e-06 *** |
|  | Serotype::Background | 1.087 | 0.384 |
| *Fitness* | Serotype | 24.580 | < 2e-16 *** |
|  | Genetic Background | 1.552 | 0.201 |
|  | Serotype::Background | 6.613 | 1.89e-10 *** |
| *Bile salts –*<br>*DCH 0.05%* | Serotype | 1.304 | 0789 |
|  | Genetic Background | 0.748 | 0.153 |
|  | Serotype::Background | 1.457 | 0.196 |
| *H2O2 – 10mM* | Serotype | 2.652 | 3.14e-02 * |
|  | Genetic Background | 15.569 | 4.74e-10 *** |
|  | Serotype::Background | 1.999 | 0.141 |
| *Biofilms* | Serotype | 18.468 | 4.16e-06 *** |
|  | Genetic Background | 41.174 | 9.57e-15 *** |
|  | Serotype::Background | 9.679 | 4.40e-07 *** |
| *Hypermucoviscosity* | Serotype | 12.771 | 7.91e-07 *** |
|  | Genetic Background | 6.137 | 2.51e-04 *** |
|  | Serotype::Background | 3.043 | 2.07e-03 ** |

**Table S2.** Statistical analyses – Stepwise regression for all traits analysed.

| **Trait** | **Step** | **AIC** |
| --- | --- | --- |
| *Capsule production* | + Genetic Background | 58.87 |
|  | + Serotype | -19.96 |
| *Growth* | + Genetic Background | -734.64 |
|  | + Media | -741.30 |
| *Growth in LB* | + Serotype | -478.05 |
|  | + Genetic Background | -485.69 |
| *Growth in M02* | + Genetic Background | -344.23 |
| *Fitness* | + Serotype | -950.65 |
| *Bile salts – DCH 0.05%* | / | / |
| *H2O2 – 10mM* | + Genetic Background | -142.912 |
| *Biofilms* | + Genetic Background | -203.560 |
|  | + Serotype | -221.601 |
| *Hypermucoviscosity* | + Serotype | -213.22 |
|  | + Genetic Background | -224.40 |

**Table S3.** Mutations identified in intermediately capsulated clones from evolving populations during adaptation to nutrient rich medium.

| Strain | Population | Capsule-related mutations | Divisiome-related mutations |
| --- | --- | --- | --- |
| *Kpn* NTUH K2044 | 1 | 9 G repeat instead of 10 – *rmpA* | Δ1,451bp [*tatC–tatD–rfaH*] |
| *Kpn* NTUH K2044 | 2 | 9 G repeat instead of 10 – *rmpA*<br>L33* – *wzc* | Δ1,073bp [*tatC–tatD–rfaH*] |
| *Kpn* NTUH K2044 | 3 | 9 G repeat instead of 10 – *rmpA* | Δ8bp – [*ftsX*] |
| *Kpn* NTUH K2044 ΔK1::K1 | 1 | IS – *wcaJ* |  |
| *Kpn* NTUH K2044 ΔK1::K1 | 2 | G151S – *wcaJ* | *354G – *zipA* |
| *Kpn* NTUH K2044 ΔK1::K1 | 3 | Δ3,307bp<br>[DNJGBDGM_03300/*galF*] | Q126* – *envC* |
| *Kpn* NTUH K2044 ΔK1::K2 | 1 | (7A repeat --> 8) +1A – *wcaJ*<br>Δ76,191bp – hvplasmid | Δ305bp [*tatC–tatD*] |
| *Kpn* NTUH K2044 ΔK1::K2 | 2 | L142R – *rfaH*<br>2bp – *ugd* | V145G – *tatC* |
| *Kpn* NTUH K2044 ΔK1::K2 | 3 | 2bp – *ugd* |  |
| *Kpn* NTUH K2044 ΔK1::K3 | 1 |  | V145G – *tatC* |
| *Kpn* NTUH K2044 ΔK1::K3 | 2 | (ACGCCG)x2 – *galE* | Δ1bp – *tatC* |
| *Kpn* NTUH K2044 ΔK1::K24 | 1 | Δ297bp – *wcaJ* |  |
| *Kpn* ST45 | 1 | Frameshift (+T) – *wcaJ* | Q90* (CAG --> TAG) – *tatC* |
| *Kpn* ST45 ΔK24::K1 | 1 |  | V145G – *tatC* |
| **Kpn ST45 ΔK24::K1** | 2 | Δ1bp – *wcaJ* | V145G – *tatC* |
| **Kpn ST45 ΔK24::K1** | 3 | Δ1bp – *wcaJ* | V145G – *tatC* |
| **Kpn ST45 ΔK24::K24** | 1 | I65N – *wcaJ* | V145G – *tatC* |
| **Kpn ST45 ΔK24::K24** | 2 |  | V145G – *tatC* |

